# CD133 shapes extracellular vesicle cargo and angiogenic function in basal-like triple-negative breast cancer

**DOI:** 10.1101/2025.10.31.685601

**Authors:** Mireia Gomez-Duro, Ptissam Bergam, Lorena Martin-Jaular, Sara Sanchez-Redondo, Marta Hergueta-Redondo, Florent Dingli, Damarys Loew, Hector Peinado, Thierry Dubois, Graça Raposo, Gisela D’Angelo

## Abstract

Extracellular vesicles (EVs) are central mediators of tumor-stroma communication, yet the mechanisms governing their biogenesis and functional cargo remain poorly understood. Here, we identify CD133 (Prominin-1), selectively expressed in basal-like triple-negative breast cancer (BL-TNBC) cells, as a key regulator of EV production and cargo composition. CD133 localizes to plasma membrane protrusions and lipid rafts, maintaining membrane architecture, promoting EV release, and shaping selective protein loading, including the pro-angiogenic factor CD105. Functionally, CD133⁺ EVs potently induce endothelial tubulogenesis in a CD105-dependent manner, independently of EV uptake or endothelial proliferation. *In vivo*, these EVs preferentially accumulate in the lung and liver, highlighting organ-specific communication. Mechanistically, CD133 maintains lipid raft integrity, facilitating CD105 selective incorporation into vesicles, thereby establishing a spatially coordinated CD133-CD105 axis. Together, our findings define a subtype-specific EV population whose identity and angiogenic activity are dictated by membrane organization. By linking protrusion architecture to vesicle-mediated signaling, this study highlights CD133-enriched EVs as potential biomarkers and therapeutic targets in aggressive BL-TNBC.

## INTRODUCTION

Triple-negative breast cancer (TNBC) is a highly aggressive subgroup of breast cancer (BC) defined by the absence of estrogen and progesterone receptor expression and *HER2* amplification. Although TNBC accounts for 15-20% of all BC (Loibl et al., 2021), it is disproportionately associated with early relapse, metastasis, and poor outcomes. This is largely due to its limited therapeutic options and intrinsic molecular heterogeneity (Bianchini et al., 2022). Transcriptomic profiling has delineated TNBC into several molecular subtypes with distinct signaling pathways, including basal-like (BL1 and BL2), mesenchymal (MES), and luminal androgen receptor (LAR), each subtype exhibiting unique biological behaviors and therapeutic responses (Lehmann et al., 2011; Lehmann et al., 2016; Echevarria et al., 2018; Bareche et al., 2018; Lehmann et al., 2021). The BL subtypes are the most prevalent and enriched for genes involved in cell cycle control, DNA damage repair, and stemness pathways. These features may influence both therapeutic resistance and communication with the tumor microenvironment (TME) (Lehmann et al., 2011; Lehmann et al., 2016; Bianchini et al., 2022).

Given the aggressive nature of TNBC and its reliance on TME interactions, extracellular vesicles (EVs) have emerged as central mediators of tumor–stromal communication, enabling cancer cells to modulate their surroundings and disseminate pro-tumorigenic signals (Peinado et al., 2012; Hoshino et al., 2015; Becker et al., 2016; Xu et al., 2018). EVs are membrane-enclosed particles that transport proteins, lipids, and nucleic acids to recipient cells, influencing cellular behavior in both local and distant tissues (van Niel et al., 2018; D’Angelo et al., 2025). Tumor-derived EVs contribute to invasion, immune evasion, therapy resistance, and the establishment of pre-metastatic niches (PMN) (Clancy & D’Souza-Schorey, 2023). Importantly, EVs are heterogeneous, comprising functionally distinct subpopulations that differ in size, origin, and cargo. These include endosome-derived exosomes, ectosomes that bud directly from the plasma membrane, and protrusion-derived EVs (PD-EVs) generated at specialized membrane extensions such as microvilli, filopodia, and cilia (van Niel et al., 2018; Marzesco et al., 2005; Dubreuil et al., 2007; Rilla et al., 2021; Nishimura et al., 2021; Hurbain et al., 2022; D’Angelo et al., 2023; Hu et al., 2024; Gomez-Duro et al., 2025). While exosomes and ectosomes have been extensively characterized, PD-EVs remain less well understood. Notably, our recent study in Drosophila demonstrated that microvilli-derived CD133⁺ EVs mediate the polarized delivery of the Hedgehog morphogen, providing direct evidence that PD-EVs can serve as instructive vehicles for directional intercellular communication (Hurbain et al., 2022).

CD133 (Prominin-1) is a pentaspan transmembrane protein localized to membrane protrusions and lipid rafts, sites associated with EV biogenesis (Weigmann et al., 1997; Röper et al., 2000; Corbeil et al., 2001). Initially described as a cancer stem cell (CSC) marker across multiple tumor types (Singh et al., 2004; O’Brien et al., 2007; Li et al., 2017; Glumac & LeBeau, 2018), emerging evidence indicates that CD133 is not merely a passive marker but actively contributes to tumor progression by influencing signaling and TME interactions (Brugnoli et al., 2019). In human cancers, CD133 expression correlates with tumor-initiating capacity, therapeutic resistance, and poor prognosis (Currie et al., 2013; Bareche et al., 2020). In breast cancer, expression is highly variable, ranging from 1–2% in some lines to nearly 80% in BL-TNBC cells such as MDA-MB-468 (Borgna et al., 2012). This variability, combined with its localization to plasma membrane protrusions and its role in EV release in Drosophila, raises the possibility that CD133 actively regulates EV production and cargo loading in human tumors.

In this study, we establish CD133 as a central organizer of EV biogenesis and function in BL-TNBC. CRISPR-Cas9-mediated CD133 knockout combined with proteomic analyses reveal that CD133 governs selective cargo loading, including incorporation of the pro-angiogenic factor CD105. CD133⁺ EVs potently stimulate endothelial tubulogenesis in a CD105-dependent manner and preferentially distribute to the lung and liver, suggesting a role in early organotropic signaling. By contrast, long-term preconditioning with either CD133⁺ or CD133 depleted-EVs similarly promotes metastatic colonization and vascular remodeling, indicating that CD133 primarily shapes the specificity of early EV-mediated communication rather than sustaining late-stage progression. Collectively, these findings uncover a CD133-CD105 axis that endows EVs with pro-angiogenic activity in BL-TNBC, redefining CD133 from a passive CSC marker to an active regulator of vesicle-mediated signaling with implications for angiogenesis, metastasis, and therapeutic targeting in this aggressive breast cancer subtype. BL-TNBC.

## MATERIAL AND METHODS

### Ethical Regulations

All research conducted in this study adhered to relevant ethical guidelines. For retrospective clinical studies, written informed consent was obtained from all patients, who either agreed or did not explicitly refuse the use of their data for research. Animal experiments were carried out following approved protocols: PROEX178/15 and PROEX225/17 (ISCIII/Comunidad de Madrid, Spain) and (DAP 2024-011/Institut Curie/Paris, France). All procedures complied with the International Guiding Principles for Biomedical Research Involving Animals (CIOMS). When applicable, tumor volumes did not exceed the maximum limit of 1500 mm³ as approved by the ISCIII Ethical Committee.

### Transcriptomic analyses

#### Curie cohort

Our cohort has been previously described and is composed of 35 luminal A (Lum A), 40 luminal B (Lum B), 46 TNBC, 33 HER2^+^, 18 normal breast tissues (HD) (Maire et al., 2013a; 2013b; Suresh et al., 2022). Experiments were conducted in agreement with the Bioethic Law No. 2004–800 and the Ethical Charter from the French National Institute of Cancer (INCa), and after approval of the ethics committee of our Institution. RNA profiling using Affymetrix U133 Plus 2.0 microarrays (ThermoFisher Scientific) on this cohort have been previously described (Maire et al., 2013a; 2013b; Suresh et al., 2022).

#### Breast (cancer) cell lines

Total RNA from the different breast (cancer) cell lines (Maire et al. 2013a; 2013b; Suresh et al., 2022) was extracted following the procedure described by Dakroub *et al*. (Dakroub et al., 2023) using the miRNeasy Kit (Qiagen, Courtaboeuf, France) following the manufacturer’s protocol. RNA concentration and purity were assessed using a NanoDrop ND-1000 spectrophotometer (Thermo Fisher Scientific), while RNA integrity was evaluated via the RNA 6000 Nano LabChip on an Agilent 2100 Bioanalyzer (Agilent Technologies). High-quality RNA samples were subsequently hybridized to GeneChip™ Human Exon 1.0 ST arrays (Affymetrix), following the manufacturer’s recommended workflow. Raw data were processed and subjected to quality control using Affymetrix Expression Console Software. Downstream analysis of the transcriptomic profiles was performed as previously described (Lerebours et al., 2021).

### Survival analyses

Kaplan-Meier curves for recurrence free survival (RFS) of breast cancer patients stratified by high and low expression of *PROM1/CD133* mRNA (probeset 204304_s_at, best cutoff setting) was determined from the online tool (https://www.kmplot.com) (Lánczky & Győrffy, 2021).

### Mice

5-7-week-old athymic nude female mice from Charles River were used unless otherwise specified for EV *in vivo* distribution, EV conditioning, and vascular leakiness. Animals were housed at 21°C +/−2°C, and humidity was 50-60%. The light cycle was light:13 hours/dark: 11 hours.

### Cell culture

Cell lines were obtained from the American Type Culture Collection (ATCC, Manassas, VA, USA). MDA-MB-231 and MDA-MB-468 cells were cultured in DMEM, high glucose, GlutaMAX™ Supplement (Thermo Fisher Scientific), supplemented with 10% FBS (Gibco), 5% Pen/Strep (Gibco). HCC-70 cells were maintained in RPMI-1640 GlutaMAX™ medium (Sigma-Aldrich) supplemented with 1.5 g/L sodium bicarbonate, 10 mM HEPES, and 1 mM sodium pyruvate (all from Thermo Fisher Scientific), 10% FBS, and 5% Pen/Strep. Modified MDA-MB-468 cell lines, including CD133 KO, scramble control and shCD105 KD cells, were maintained in complete DMEM medium under puromycin selection (2 µg/mL, Thermo Fisher). Primary HUVECs were purchased from Promocell (C-12203) and cultured in Endothelial Cell Growth Medium 2 Supplement (PromoCell) supplemented with 0.02 mL/mL FCS, 5 ng/mL recombinant human EGF, 10 ng/mL recombinant human bFGF, 20 ng/mL recombinant human long R3 IGF, 0.5 ng/mL recombinant human VEGF165, 1 µg/mL ascorbic acid, 22.5 µg/mL heparin, and 0.2 µg/mL hydrocortisone, according to the manufacturer’s instructions.

All cells were incubated at 37°C in a humidified atmosphere with 5% CO₂. Upon acquisition, cells were expanded and frozen in aliquots stored in vapor-phase liquid nitrogen. Cell stocks were used up to passage 20 and discarded thereafter, except for HUVECs, which were used only up to passage 5. All cultures tested negative for Mycoplasma by PCR.

### Antibodies

The following antibodies were used for WB, IF, or IEM: rabbit (rb) anti-β3-tubulin (Invitrogen 2G10MA1-118; 1:500 WB); rb anti-GAPDH (Sigma G9545, 1:10.000 WB); ms anti-β-actin (Sigma A2228; 1 :10.000 WB); rb Syntenin (Abcam ab133267; 1:1.000 WB) or (Abcam ab19903; 1:1.000 WB); rb anti-Alix (Abcam ab186429; 1:1.000 WB); ms anti-Alix (Cell Signaling 2171S ; 1:1.000 WB); ms monoclonal anti-TSG101 (Santa Cruz, C-2, sc-7964; 1:200 WB); ms anti-human CD81 monoclonal antibody (Millipore CLB579; 1:1.000 WB) or (Santa Cruz sc-166029; 1:50 WB); ms anti-CD9 (Sigma-Aldrich, clone MM2/57 CLB 162; 1:1.000 WB); ms monoclonal CD63 (Invitrogen, 1:200 WB; 1:50, IF), (Thermo Fisher Scientific MA1-35272; 1:200 WB; 1:50 IEM, IF) or (BD Pharmingen 556019; 1 :1.000 WB); ms monoclonal anti-TSG101 (Santa Cruz, C-2, sc-7964; 1:200 WB); ms anti-flotilin-1 (BD Biosciences 610821; 1:1.000 WB); rb CD9 (Abcam ab236630; 1:80 IEM); rb CD81 (Abcam ab233692; 1:50 IEM); rb anti-CD133 clone CD133 D2V8Q (Cell Signaling 64326; 1:1.000 WB; 1:200 IF); rb anti-CD133 D4W4N XP (Cell Signaling 86781; 1 :700 IHC); anti-CD133/1 (Miltenyi Biotec 130-113-670; 1:1.000 WB, 1:50 IEM); PE monoclonal anti-human CD133/1 clone AC133 (Miltenyi Biotec; 1:50 FC); ms anti-Endoglin (Santa Cruz sc-18838; 1:10 IF, IEM); rb anti-endoglin (Proteintech 10862-1-AP; 1:1.000 WB; 1:100 IP, IF); rb anti-IgG isotype control (Invitrogen; co-IP); ms monoclonal anti-Pan Cytokeratin Monoclonal Antibody (AE1/AE3) (Thermo Fisher Scientific 53-9003-82; 1 :500 IHC) and . Secondary antibodies coupled to HRP-conjugated goat polyclonal antibodies to rabbit IgG (Abcam ab6721) or to mouse IgG (Abcam ab6789) were used at 1:5,000. Secondary goat anti-rabbit or anti-mouse antibodies conjugated to Alexa Fluor 488, 555, or 647 were from Invitrogen and used at 1:200. Alexa Fluor™ 647 Phalloidin (Thermo Fisher, A22287).

### Generation of CD133 KO and CD105 KD MDA-MB-468 Cells

#### CRISPR/Cas9-mediated CD133 knockout (CD133 KO)

CD133 KO in MDA-MB-468 cells was performed using the dual-guide lentiviral plasmid pLV[2CRISPR]-hCas9 (VectorBuilder, VB221117-1157xvx), which encodes two gRNAs targeting the PROM1 gene (gRNA1: GTGGCGTGTGCGGCTATGAC; gRNA2: GTTTGGCGTTGTACTCTGTC), hCas9, and a puromycin resistance cassette. A scramble control vector carrying non-targeting gRNAs was used in parallel. Plasmids were propagated in E. coli and purified using EndoFree Midiprep kits before lentiviral production.

Lentiviral particles were generated by co-transfecting HEK 293-LTV packaging cells with the CRISPR/Cas9 vector, the packaging plasmid psPAX2, and the envelope plasmid pCMV-VSV-G, using TransIT®-293 Transfection Reagent (Mirus Bio) according to the manufacturer’s instructions. Viral supernatants were collected at 48 h, filtered (0.22 µm), and concentrated by ultracentrifugation at 120,000 × g for 90 min (SW32 Ti rotor; Beckman Coulter). MDA-MB-468 cells were transduced with the viral supernatants and selected with puromycin (2 µg/mL) for 7 days. CD133 KO efficiency was confirmed by sequencing, RT-qPCR, and flow cytometry.

#### shRNA-mediated CD105 knockdown (KD)

For CD105 KD, MISSION® TRC2 pLKO.5-puro lentiviral vectors (Sigma-Aldrich) were used, including a non-targeting control shRNA (SHC202) and two CD105-targeting shRNAs: TRCN0000003273 (shCD105#1) and TRCN0000003276 (shCD105#2). The resulting constructs, shCD105#1 or shCD105#2, were verified by sequencing.

Lentiviral particles were produced by transfecting HEK-Lenti-X cells with 0.5 µg of each shRNA plasmid, 0.375 µg psPAX2, and 0.125 µg pMD2.G per well using Lipofectamine P2000 (ThermoFisher, Cat. 11668027). After 2 h at 37°C, 1 mL of DMEM without antibiotics or FBS was added. Sixteen hours later, the medium was replaced with DMEM containing 30% FBS. Viral supernatants were collected over 2 days, clarified by centrifugation (2,000 × g, 10 min, 4°C), and used to infect MDA-MB-468 cells (3 × 10⁶ cells per 100 mm plate). Infected cells were selected with puromycin (2 µg/mL) for at least 120 h, and knockdown efficiency was confirmed by Western blot. Puromycin was maintained throughout to ensure KD stability.

#### Quantitative real-time PCR (qPCR)

Total RNA was extracted from MDA-MB-468 (WT), CD133 KO, or scramble control cells using the NucleoSpin RNA extraction kit (MACHEREY-NAGEL), according to the manufacturer’s instructions. Complementary DNA (cDNA) was synthesized from 0.3 µg of total RNA using the SuperScript First-Strand Synthesis System (Invitrogen; catalog no. 11904-018). Primers for CD133 (forward 5′-GGCCCAGTACAACACTACCAA-3′; reverse 5′-ATTCCGCCTCCTAGCACTGAA-3′) were selected from PrimerBank (ID: 224994190c2) after in silico validation of GC content, melting temperature, amplicon size (75 bp), and absence of secondary structures. qPCR was performed using the LightCycler 480 system (Roche), with GAPDH as the endogenous reference gene (forward 5′-CTGGGCTACACTGAGCACC-3′; reverse 5′-AAGTGGTCGTTGAGGGCAATG-3′). Each reaction was run in triplicate across at least three biological replicates, and relative gene expression was calculated using the 2ΔΔCT method. Knock-out clones were further validated by sequencing to confirm the targeted deletion.

### Surface biotinylation assay

WT and KO CD133 cells were grown to ∼ 80% confluence in four 150 mm culture dishes. Cells were incubated in Opti-MEM for 30 min at 37°C, then detached with PBS-2mM EDTA and washed with ice-cold PBS. Surface proteins were biotinylated by incubating cells with 0.2 mg/mL of EZ-Link Sulfo-NHS-LC-Biotin (Thermo Fisher Scientific; catalog no. 21335) in cold PBS for 30 min at 4°C. Following biotinylation, cells were washed twice with ice-cold PBS and quenched by incubation with 50 mM NH_4_Cl for 3 min at 4°C to inactivate the unreacted biotinylation reagent. Cells were then washed three times with ice-cold PBS and pelleted by centrifugation at 200*g* for 5 min at 4°C. Cell pellets were lysed in a homemade RIPA buffer composed of 50 mM Tris-HCl (pH 7.4), 150 mM NaCl, 1% NP-40, 0.5% sodium deoxycholate, 0.1% SDS, and 1 mM EDTA, supplemented with a protease inhibitor cocktail (Sigma-Aldrich; P8340). Lysates were incubated on ice for 20 minutes at 4°C to ensure complete lysis. Lysates were centrifuged at 20,000*g* for 20 min at 4°C, and the resulting supernatant was collected. Protein concentrations were quantified using BioRad Protein Assay reagent (BioRad), and equal amounts of total protein were incubated with 100 μl of prewashed NeutrAvidin Plus UltraLink Resin (Thermo Fisher Scientific; catalog no. 53151) for 3 h on an orbital shaker at 4°C. Beads were washed five times with 1 mL of cold RIPA buffer, and bound proteins were eluted from beads with 30 μl of Laemmli sample buffer (BioRad; catalog no. 1610747). Eluted proteins and 1% of the input lysate (total protein) were analyzed by WB analysis to assess the surface expression of specific proteins.

#### Lipid raft enrichment (by Detergent-Resistant Membrane Fractionation)

Lipid raft-enriched membrane fractions were isolated from WT and CD133 KO cells following an adapted protocol described by da Silva-Januario *et al*. (da Silva-Januario et al., 2023) 5.0 × 10^7^ were lysed in 1 mL of ice-cold TNE buffer (10 mM Tris–HCl, pH 7.5, 150 mM NaCl, 5 mM EDTA) containing 0.5% Triton X-100, a protease inhibitor cocktail (Sigma-Aldrich; P8340) and a phosphatase inhibitor cocktail (Sigma-Aldrich; catalog no. P0001). Lysates were passed 20 times through a 21-gauge needle and centrifuged at 1,000*g* for 10 min at 4°C, and a total of 950 μL of the resulting supernatant was layered on the bottom of a 16 mL polypropylene ultracentrifuge tube (Beckman Coulter; 326823). A discontinuous sucrose gradient was formed by sequentially overlaying the lysate with 4 mL of 85% sucrose in TNE buffer, followed by 5 mL of 35% sucrose, and 5 mL of 5% sucrose, all in TNE buffer. Samples were ultracentrifuged in a SW32.1 Ti -16 mL rotor (Beckman Coulter) at 155,846 *g* for 24h at 4°C. Fractions of 1.6 mL were collected from the top of the gradient, and proteins in each fraction were precipitated overnight at 4°C with 15% TCA (JT Baker; catalog no. 0414-04), washed once with cold acetone, and air-dried at RT. Pellets were resuspended in 20 µl of Laemmli sample buffer containing 100 mM DTT (BioRad; catalog no. 1610747), boiled at 95°C for 5 min, and submitted to a 4-15% Mini-protean TGX Gel (BioRad; catalog no. 4561083), followed by WB analysis.

### Serum EV-depleted medium

To eliminate EVs from FBS, 250 mL of RPMI or DMEM containing 20% FBS was ultracentrifuged at 100,000 × g for 18 hours at 4°C using a Ti45 rotor (Beckman Coulter). After ultracentrifugation, the EV-depleted supernatant was collected and supplemented with 5% Pen/Strep and 250 mL of fresh RPMI or DMEM to obtain the final working concentration. The mixture was then filtered through a 0.22 µm Stericup sterile vacuum filtration system (Millipore Sigma). This FBS-EVs-depleted medium was used in all EV-related experiments, except in those involving proteomic analysis, where commercially available exosome-depleted FBS (Thermo Fisher Scientific) was used.

### EV isolation

EVs were isolated from TNBC cell-conditioned medium (CM) using either differential ultracentrifugation (dUC) or size exclusion chromatography (SEC). For both methods, cells were washed with PBS and reincubated with DMEM or RPMI supplemented with 10% exosome-depleted FBS (Thermo Fisher Scientific, A2720801) or in serum-EVs-depleted medium (prepared in-house as described above). For EV isolation, CM were collected after 24 hours from TNBC cell cultures at 70– 90% confluence. Cells were counted and viability was assessed before harvesting. EV concentration, size distribution, and morphology were determined by NTA, nFCM, WB, TEM, IEM, and surface marker profiling via bead-based multiplex FC with the MACSPlex EV Kit IO (Miltenyi Biotec). All EV isolation procedures followed the MISEV2018 and MISEV2023 guidelines (Théry et al., 2018; Welsh et al., 2023).

#### By differential ultracentrifugation (dUC)

EVs were isolated by dUC following a standard protocol (Théry et al., 2006). CM was subjected to a series of centrifugation steps: first at 300 × g for 10 minutes at 4°C to remove cells, followed by 2,000 × g for 20 minutes at 4°C to eliminate cellular debris, and then at 10,000 × g for 30 minutes at 4°C to remove larger vesicles and apoptotic bodies. The resulting supernatant was then ultracentrifuged at 100,000 × g for 60 minutes at 4°C using a Ti45 rotor (Beckman Coulter; catalog no. 355622) to pellet the small EVs. The pellet was resuspended in PBS (pH 7.5), washed, and centrifuged again at 100,000 × g for 60 minutes at 4°C using a Ti70 rotor (Beckman Coulter; catalog no. 355618). Final EV pellets were resuspended in PBS and either stored at –80°C or used immediately, or after overnight storage at 4°C in non-frozen conditions.

#### By size exclusion chromatography (SEC)

CM was harvested by pelleting cells at 300 × g centrifugation, at 4°C. The supernatant was further centrifuged at 2,000 × g for 20 minutes at 4°C to remove cell debris. The resulting supernatant was concentrated using Centricon Plus-70 centrifugal filters (Millipore, 10 kDa molecular weight cutoff) at 2,000 × g at 4°C until the final volume was reduced to less than 500 µL. The concentrated CM was then applied to qEV original SEC columns (Izon, SP5), which allowed optimal recovery of vesicles ranging from 35 to 350 nm. The void volume (fractions 1–6, totaling 3 mL) was discarded, and fractions 7-11 (2.5 mL), enriched in EV, were collected. These fractions were then concentrated using Amicon Ultra-15 centrifugal filters (Millipore, 10 kDa cutoff) at 3,320 × g at 4°C.

Aliquots of the final EV preparations were stored at −80°C until use. In some cases, vesicles were stored overnight at 4°C and used without freezing.

### Single-particle characterization of EVs

#### Nanoparticle Tracking Analysis (NTA)

NTA was performed using the ZetaView PMX-120 system (Particle Metrix, software v8.04.02) at 22°C. Instrument settings were sensitivity 75, shutter speed 75, and frame rate 30 fps. Two independent dilutions of each sample were analyzed across 11 positions, with two cycles per position.

#### Nano-Flow Cytometry (nFCM)

For high-resolution single-particle analysis, nFCM was carried out using the NanoAnalyzer (NanoFCM, S/N FNAN30E22101948). The instrument used single-photon-counting modules to detect side scatter (SSC, 488/10 nm), green fluorescence (525/40 nm), and far-red fluorescence (670/30 nm). Calibration for particle concentration was done using 250 nm fluorescent silica QC beads, and particle sizing was calibrated with a standard mix of silica beads (68, 91, 113, and 155 nm; S16M-Exo beads). Samples were diluted in 0.2 µm-filtered 1× PBS to reach a final concentration of 2,000–12,000 particles/min, measured at 1 kPa pressure. Data were recorded using NanoFCM Professional 2.0 software and exported as .fcs files for analysis in FlowJo v10.10.0.

### Multiplex bead-based flow cytometry for EV surface marker profiling

EVs isolated from TNBC cells via SEC were characterized using bead-based multiplex FC (MACSPlex EVs Kit IO, human; Miltenyi Biotec). EV input was standardized based on particle concentration determined by NTA. A total of 1 × 10⁹ EVs were diluted in MACSPlex buffer to a final volume of 120 µL and incubated overnight at RT with 10 µL of MACSPlex Exosome Capture Beads on an orbital shaker, protected from light. Following incubation, samples were washed and stained for 1 hour at RT with APC-conjugated detection antibodies against CD9, CD63, and CD81 (provided in the kit), either individually or in combination. Flow cytometric analysis was performed using a Spectral Flow Cytometer Analyzer Northern Lights (Cytek Biosciences) according to the manufacturer’s recommendations, and data were analyzed with FlowJo software (v10, FlowJo LLC). Median fluorescence intensity (MFI) values were background-corrected by dividing the signal by the fluorescence of capture beads incubated with detection antibodies in the absence of EVs. Among the 37 markers included in the assay, only selected markers relevant to the study objectives are reported.

### Western Blot analysis (WB)

Cells were harvested and washed twice with cold PBS. Approximately 1 × 10⁷ cells were pelleted by centrifugation at 300 × g for 5 minutes at 4°C and resuspended in 200 µL of lysis buffer containing 50 mM Tris-HCl (pH 7.5), 150 mM NaCl, 10% (v/v) glycerol, 5 mM EDTA, and 1% (v/v) Triton X-100, supplemented with a protease inhibitor cocktail (Sigma-Aldrich; P8340). Lysates were incubated on ice for 20 minutes with gentle agitation, followed by centrifugation at 16,000 × g for 20 minutes at 4°C to remove insoluble material. The resulting clarified supernatants were quantified using the Pierce™ BCA Protein Assay Kit (Thermo Fisher Scientific; catalog no. 23225). Both whole-cell lysates (12 µg of protein per lane) and EVs (1 × 10⁹ particles) were mixed with Laemmli sample buffer (BioRad; catalog no. 1610747), with or without 400 mM DTT (BioRad; catalog no. 1610747), and denatured by boiling for 5 minutes at 95°C or for 10 minutes at 70°C, respectively. Proteins were separated on NuPAGE (4-12%) Bis-Tris gels (Invitrogen) and transferred onto nitrocellulose membranes (Millipore). Membranes were blocked for 1 hour at RT in PBS/ Tween-20 0.1% (PBS/T) with 5% non-fat dry milk, incubated overnight at 4°C with the indicated primary antibodies diluted in the same blocking solution. After three washes with PBS/T, membranes were incubated with HRP-conjugated secondary antibodies, followed by additional washes in PBS/T. Immunoreactive bands were visualized using Clarity™ Western ECL SubstrateBio-Radad, catalog no. 170-5060) and detected using the ChemiDoc Touch Imaging System (Bio-Rad), ensuring exposure within the linear detection range. Band intensities were quantified by densitometry using the “Analyze Gels” tool in ImageJ (https://imagej.net/ij/).

### Immunofluorescence (IF)

WT and CD133 KO cells were seeded in μ-Slide 8-well chamber slides (Ibidi; catalog no. 80826) and cultured for 24 hours at 37°C in a humidified atmosphere containing 5% CO₂. For surface immunolabeling protocol was adapted from Thamm *et al*. (Thamm et al., 2019). Cells were gently rinsed with calcium/magnesium buffer (PBS supplemented with 1 mM CaCl₂ and 0.5 mM MgCl₂) and incubated in blocking buffer (Ca/Mg buffer with 0.2% gelatin) for 30 minutes at 4°C. Cells were then incubated with primary antibodies diluted in a blocking buffer for 1 hour at 4°C. In selected experiments, after labeling for CD133, cells were incubated with rhodamine-conjugated WGA (10 µg/mL; Vector Laboratories) for 30 minutes at 4°C. Following antibody or WGA incubation, cells were washed with PBS and fixed in 4% PFA (Electron Microscopy Sciences; catalog no. 15710) in PBS for 30 minutes at RT. After fixation, cells were quenched for 10 minutes in PBS containing 50 mM NH₄Cl to reduce background fluorescence. Secondary antibodies, diluted in a blocking buffer, were added for 1 hour at RT. In some experiments, nuclei were counterstained with DAPI (1 µg/mL; Sigma-Aldrich) for 10 minutes at RT. After staining, samples were washed three times with PBS containing 0.2% gelatin, followed by two washes in PBS and a final rinse in distilled water. Cells were mounted using Mowiol 4.88 (Merck) or Fluoromount-G Mounting Medium (Electron Microscopy Sciences; catalog no. 17984-25).

### Image acquisition, analysis, and quantification

Confocal imaging was performed using a Zeiss LSM 780 confocal laser scanning microscope (Zeiss, Jena, Germany) equipped with a 63× oil immersion objective (NA 1.4). Z-stack images were acquired at an interval of 0.5 µm or less under identical laser, gain, and offset settings across all experimental conditions. Post-acquisition image processing and quantification were conducted using ImageJ software (version 1.36; NIH). Co-localization analysis was performed using Z-stack image sets from individual cells, and Mander’s overlap coefficients (tM) were calculated using the Colocalization Threshold plugin in ImageJ (Thamm et al., 2019).

### Flow cytometry (FC) of CD133 expression

WT, scramble control and CD133 KO MDA-MB-468 cells were harvested using 0.05% trypsin/0.5 mM EDTA solution (Life Technologies) for 5 minutes at 37°C. Cells were centrifuged, and the resulting pellet was resuspended in PBS containing 5 mM EDTA (VWR International). Cell suspensions containing 2 × 10⁵ cells in 100 µL were incubated with or without PE-conjugated monoclonal antibody against human CD133 for 30 minutes at 4°C. After incubation, cells were washed with PBS and resuspended in PBS with 5 mM EDTA. DAPI was added before acquisition to exclude non-viable cells. Instrument settings and gating strategies were determined using appropriate controls, including unstained and CD133-stained populations. During analysis, viable cells (DAPI–) were first selected, followed by singlet discrimination using FSC-A versus FSC-H to exclude doublets. CD133 expression was quantified within this gated live singlet population. A total of 20,000 events per sample were acquired using a FACSCanto II flow cytometer (BD Biosciences), and data were analyzed using FlowJo software (v10, FlowJo LLC).

### Transmission Electron Microscopy (TEM)

Ultrastructural analyses of cells and EV samples were performed according to standard protocols and MISEV guidelines (Théry et al., 2018).

#### Cellular TEM (chemical fixation)

TNBC cell monolayers were fixed in a solution of 2.5% glutaraldehyde (Electron Microscopy Sciences; catalog no. 16220) and 2% PFA (Electron Microscopy Sciences; catalog no. 15710) in 0.1 M cacodylate buffer for 24 hours at 4°C. Post-fixation was performed using 1% osmium tetroxide supplemented with 1.5% potassium ferrocyanide, followed by graded ethanol dehydration and embedding in EPON resin. Ultrathin sections (60 nm) were cut, mounted on grids, and contrasted with uranyl acetate (UA) and lead citrate.

#### Cellular TEM (High-Pressure Freezing (HPF) and Freeze Substitution)

HPF was employed to preserve cellular ultrastructure in a near-native state by preventing ice crystal formation and associated artifacts. Cells were cultured on sapphire discs, selected for their thermal conductivity and durability, and rapidly vitrified under ∼2000 bar (200 MPa) pressure using the HPM CryoCapCell system with liquid nitrogen cooling. Ethanol injections stabilized temperature and pressure between cycles. Frozen samples were transferred to cryogenic tubes and subjected to automated freeze substitution (AFS), during which 1% osmium tetroxide was added, and the temperature was gradually increased from −90°C to 0°C over several days. Post-substitution, samples were mounted on glass slides, fixed in 2 mM sodium cacodylate buffer, rinsed with PBS, and dehydrated through graded ethanol baths (50–100%) at RT. Gradual infiltration with EPON resin was performed (30% for 30 minutes, 70% for 2 hours, 100% overnight), followed by polymerization at 60°C for 48 hours. Resin blocks were trimmed with a Leica ultramicrotome into pyramidal shapes and sectioned perpendicularly with a diamond knife into 70 nm ultrathin sections. Sections were collected on 200-mesh copper/palladium grids, contrasted by UA (20 minutes, dark) and lead citrate (2 minutes, CO₂-controlled), then rinsed and stored in dust-free capsules until TEM analysis. Imaging was performed using a Tecnai Spirit G2 transmission electron microscope (ThermoFisher Scientific) operated at 80 kV, equipped with a 4k CCD camera (Gatan OnView 4k x 4k). Image acquisition and processing utilized standard digital imaging systems.

#### EV TEM

1 x 10^10^ particles quantified by NTA and resuspended in PBS (final volume of 5 µL) were deposited onto Formvar-carbon-coated EM grids and allowed to adsorb for 20 minutes at RT. Grids were fixed in 2% PFA (Electron Microscopy Sciences; catalog no. 15710) in 0.2 M PB (pH 7.4) for 20 minutes, washed six times with distilled water, and post-fixed in 1% glutaraldehyde (Electron Microscopy Sciences; catalog no. 16220) for 5 minutes. Negative staining was carried out using a mixture of 4% UA and 2% methylcellulose (1:9 ratio) for 10 minutes on ice. Excess stain was removed by blotting with Whatman paper, and the grids were air-dried for 30 minutes at RT. All staining steps were performed in the dark.

Both cellular and EV TEM samples were examined using a Tecnai Spirit G2 transmission electron microscope (Thermo Fisher Scientific) operated at 80 kV and equipped with a 4k CCD camera (On View 4k x 4k GATAN). Image acquisition and analysis were performed using iTEM software (EMSIS). Quantitative analyses and statistical processing were carried out using GraphPad Prism.

### Immunolabeling Electron Microscopy (IEM)

Immunogold labeling of cells and EVs was performed using established protocols adapted from Hurbain et al. (Hurbain et al., 2008; Posthuma, 2008).

#### Cellular IEM

Ultrathin cryosections of fixed cells were prepared using an ultracryomicrotome (Leica UC7 FCS) following fixation in 2% PFA (Electron Microscopy Sciences; catalog no. 15710) in 0.1 M PB (pH 7.4). Sections were immunolabeled using primary antibodies followed by single immunogold detection with protein A conjugated to 10 nm gold particles (Cell Microscopy Center, Department of Cell Biology, Utrecht University).

#### EV IEM

1 x 10^10^ particles quantified by NTA and resuspended in PBS (final volume of 5 µL) were deposited onto EM grids and fixed in 2% PFA (Electron Microscopy Sciences; catalog no. 15710) in 0.2 M PB (pH 7.4) for 20 minutes at RT. Immunodetection was performed using primary antibodies against CD63 (mouse monoclonal), CD9 (rabbit), or CD81 (rabbit), each diluted in 1% BSA/PBS and incubated for 1 hour at RT. For CD63, a secondary incubation with rabbit anti-mouse Fc (Dako Agilent Z0412; 1:200) was carried out for 20 minutes. All grids were then incubated with 10 nm protein A-gold (1:50 in 1% BSA/PBS), followed by postfixation with 1% glutaraldehyde (Electron Microscopy Sciences; catalog no. 16220) in PBS. Samples were negatively stained using UA and methylcellulose as described for conventional TEM.

#### Tokuyasu Immunolabeling for TEM

For immunolabeling of cellular proteins with preserved antigenicity, the Tokuyasu method followed the protocol from the Cell Microscopy Center at Utrecht Medical University (Posthuma, 2008) was used.

TNBC cells at over 80% confluence were lightly fixed overnight at 4°C in 0.1 M PB containing 0.25% glutaraldehyde (Electron Microscopy Sciences; catalog no. 16220) and 2% PFA (Electron Microscopy Sciences; catalog no. 15710) to preserve protein epitopes. After fixation, cells were washed three times with PBS and incubated for 10 minutes in PBS with 0.15% glycine to neutralize free aldehydes. Cells were embedded in gelatin by scraping, centrifugation, and resuspension in 12% gelatin, followed by incubation at 37°C and further centrifugation. Pellets were solidified on ice, trimmed, and cryoprotected overnight at 4°C in 2.3 M sucrose. Blocks were mounted on steel pins, coated with sucrose, excess removed, and rapidly frozen in liquid nitrogen. Ultrathin sections (∼90 nm) were prepared on a Leica cryo-ultramicrotome at −144°C, collected on formvar- and carbon-coated grids, and stored at 4°C. Immunolabeling was performed by incubating grids in 2% gelatin, washing with PBS-glycine, blocking with 1% BSA, and applying primary antibodies diluted in PBS/1% BSA for 1 hour at RT. After washing, mouse primary antibodies were either bridged with anti-mouse IgG or directly labeled using 15 nm colloidal gold-conjugated secondary antibodies; otherwise, grids were incubated with 10 nm protein A-gold. Following final washes, grids were fixed in 1% glutaraldehyde (Electron Microscopy Sciences; catalog no. 16220), rinsed, and contrast-stained with 2% UA and methylcellulose at pH 4 before air drying.

Both cell and EV samples were examined using a Tecnai Spirit G2 transmission electron microscope (ThermoFisher Scientific) operated at 80 kV and equipped with a 4k CCD camera (On View 4k x 4k GATAN). Image acquisition and processing were performed using standard digital imaging systems.

### Proteomics and mass spectrometry analysis

#### Sample Preparation

Frozen EVs from five biological replicates of WT and CD133 KO cells were used for proteomic analysis. Each frozen EV sample was dried and resuspended in 10 µL (2 µg/µL) of 8M urea and 200 mM ammonium bicarbonate. Reduction was performed by adding 5 mM dithiothreitol (pH 8) by vortexing for 1 h at 37°C. After cooling to room temperature, cysteines were alkylated by adding 10 mM iodoacetamide for 30 min in the dark. Samples were then diluted to 1 M urea using 100 mM ammonium bicarbonate (pH 8.0) and digested overnight at 37°C with 0.2µg trypsin/LysC (Promega). Samples were then loaded onto homemade C18 StageTips for desalting. Peptides were eluted using 40/60 MeCN/H2O + 0.1% formic acid, vacuum concentrated to dryness and reconstituted in 10µl injection buffer before liquid chromatography-tandem mass spectrometry (LC-MS/MS) analysis.

#### LC-MS/MS Analysis

Online LC was performed with an RSLCnano system (Ultimate 3000, Thermo Scientific) coupled to an Orbitrap Exploris 480 mass spectrometer (Thermo Scientific). Peptides were first trapped on a C18 column (75 μm inner diameter × 2 cm; nanoViper Acclaim PepMapTM 100, Thermo Scientific) with buffer A (2/98 MeCN/H2O in 0.1% [vol/vol] formic acid) at a flow rate of 4 µl/min over 4 min. Separation was then performed on a 50 cm x 75 μm C18 column (nanoViper Acclaim PepMapTM RSLC, 2 μm, 100Å, Thermo Scientific) regulated to a temperature of 50°C with a linear gradient of 2% to 30% buffer B (100% MeCN in 0.1% formic acid) at a flow rate of 300 nL/min over 91 min. MS full scans were performed in the ultrahigh-field Orbitrap mass analyzer in ranges m/z 375–1500 with a resolution of 120,000 at m/z 200. The top 20 most intense ions were isolated and subjected to further fragmentation via high-energy collision dissociation (HCD) activation and acquired at a resolution of 15,000 with the auto gain control target set to 100%. We selected ions with charge states from 2+ to 6+ for screening. Normalized collision energy was set at 30% and the dynamic exclusion at 40s.

#### Data Processing

For identification, the data were searched against the Homo Sapiens (UP000005640) UniProt database using Sequest HT through Proteome Discoverer (version 2.4). Enzyme specificity was set to trypsin, and a maximum of two missed cleavage sites was allowed. Oxidized methionine, Met-loss, Met-loss-Acetyl, and N-terminal acetylation were set as variable modifications. Carbamidomethylation of cysteines was set as a fixed modification. Maximum allowed mass deviation was set to 10 ppm for monoisotopic precursor ions and 0.02 Da for MS/MS peaks. The resulting files were further processed using myProMS v3.10 (https://github.com/bioinfo-pf-curie/myproms) (Poullet, Carpentier, & Barillot, 2007). FDR calculation used Percolator (The et al., 2016) and was set to 1% at the peptide level for the whole study. The label-free quantification was performed by peptide Extracted Ion Chromatograms (XICs), reextracted under conditions, and computed with MassChroQ version 2.2.21 (Valot et al., 2011).

For protein quantification, ion XICs from proteotypic peptides shared between compared conditions (TopN matching) were used with missed cleavages allowed. Median and scale normalization at the peptide level was applied on the total signal to correct the XICs for each biological replicate (n = 5) to account for sample load biases. To evaluate the statistical significance of the change in protein abundance, a linear model (adjusted for peptides and biological replicates) was performed, and a two-sided t-test was applied to the fold change estimated by the model. The p-values were then adjusted for multiple testing using the Benjamini–Hochberg FDR procedure. Proteins with at least two distinct peptides across three biological replicates, a two-fold enrichment, and an adjusted p-value ≤ 0.05 were considered in sample comparison. Proteins uniquely identified in a specific condition were included in the analysis if they met the established peptide identification criteria.

Label-free quantification (LFQ) was also performed following the algorithm as described (Cox et al., 2014), with the minimum number of peptide ratios set to 2 by allowing all peptides and with a large ratio stabilization feature. Protein values were further normalized (median & scale) to correct for potential remaining total intensity biases.

Differentially expressed proteins were defined by log₂FC thresholds and visualized in a volcano plot, including proteins uniquely detected and annotated as exclusively expressed in each condition. Hierarchical clustering analysis was conducted using z-score-transformed log₂FC and LFQ intensity values. Clustering was performed based on Euclidean distance and single linkage methods, implemented in R (https://www.r-project.org/). For heatmap visualization, LFQ intensity values were first log₂-transformed, and Z-score normalized across samples for each protein. The mean Z-score was then calculated per condition, and the 150 proteins with the highest absolute mean Z-scores were selected as representative of the EV proteomic signature (Fig. 3A, B). Gene Ontology (GO) enrichment analysis was carried out using the Enrichr platform (https://maayanlab.cloud/Enrichr/) to identify overrepresented biological processes among the 20 proteins with the highest and lowest ratios in the differential analysis of EVs derived from WT and CD133 KO cells, respectively. To investigate PPI networks centered around CD133, data were retrieved from the STRING database (https://string-db.org/) and visualized accordingly (Fig. 3C-E). All data visualizations were performed using R.

The mass spectrometry proteomics raw data have been deposited to the ProteomeXchange Consortium via the PRIDE (Perez-Riverol et al., 2025) partner repository with the dataset identifier PXD065555 (Username; Password: oQtCAsYbffyl).

### Endothelial cell tube formation assay

HUVECs were seeded at 1.9 × 10⁵ cells per well in 6-well plates and pretreated for 24 hours with EVs (6 µg/mL), VEGF (10 ng/mL), PBS (vehicle control), or left untreated. Treatments were performed in EV-depleted HUVEC medium containing 2.5% FCS. After treatment, cells were gently detached using Accutase® solution (Sigma; catalog no. A6964), counted, and seeded into pre-chilled 24-well plates coated with 250 µL of reduced growth factor Matrigel (Corning; catalog no. 356221), previously polymerized for 1 hour at 37°C. After 16-18 hours of incubation at 37°C, capillary-like tube formation was visualized using an inverted phase-contrast microscope. Quantitative analysis was conducted by capturing eight random fields per condition and measuring total branching length and junction number using the Angiogenesis Analyzer plugin for ImageJ, providing robust metrics of angiogenic network complexity.

### Cell viability and proliferation assay

To assess the biological effects of EVs from WT or CD133 KO cells on endothelial function, HUVECs were treated with the respective EVs (2.5 µg/mL) or the same volume of PBS as control and subsequently subjected to viability and proliferation assays. Cell metabolic activity was measured using the CellTiter 96® AQueous One Solution Cell Proliferation Assay (Promega), according to the manufacturer’s instructions. HUVECs (10,000 cells/well) were seeded in 96-well plates in triplicate and treated with EVs or left untreated (control). Cells were incubated for 0, 24, 48, or 72 hours. At each time point, 20 µL of reagent was added to 100 µL of culture medium per well, followed by a 2-hour incubation at 37°C with 5% CO₂. Absorbance was recorded at 490 nm using a Fluostar Optima microplate reader (BMG LabTech, Germany) to determine cell viability and proliferation.

### Assessment of EV binding and internalization in HUVECs

To assess whether CD133 affects the binding and uptake of EVs by endothelial cells, a time-course internalization assay was performed using EVs freshly isolated from WT or CD133 KO cells. EVs were labeled with 1 µl of DiD far-red fluorescent dye (Thermo Fisher Scientific) per 1 mL PBS and thoroughly washed to remove excess dye. HUVECs were seeded in 6-well plates and treated with 2.5 µg/mL of DiD-labeled EVs or left untreated, followed by incubation at 37 °C for 0, 1, 2, 3, 4, 16, or 24 hours. At each time point, EV uptake was halted by two washes with ice-cold PBS at 4°C, and cells were resuspended in PBS containing 5 mM EDTA and kept on ice. Samples were analyzed by FC using a FACSCanto II flow cytometer (BD Biosciences), acquiring 20,000 events per sample. Controls included cells incubated with PBS processed in the same way as EVs (labeled and washed), unstained cells with DAPI to assess viability, and DiD-labeled EVs alone as well as unlabeled EVs to define background signal. Gating strategies and instrument settings were established using these controls.

### Biodistribution assay

For *in vivo* experiments, CD133 KO cells were compared to scramble control cells rather than WT cells. This decision was made to minimize variability, as both CD133 KO and scramble control cells underwent the same CRISPR-Cas9 editing process, ensuring greater experimental consistency. Although scramble control cells were previously validated and shown to be comparable to WT cells, and are used here for their technical suitability.

To assess EV biodistribution, 5 μg of EVs from scramble control or CD133 KO MDA-MB-468 cells freshly isolated were labeled with 3 µl of NIR-815 infrared fluorescent dye (eBiosciences) in 1 mL of PBS. Labeled EVs were washed twice in 20 mL of PBS, collected by ultracentrifugation, and resuspended in 100 μL of PBS. The 100 µl suspension of NIR-815-labeled EVs was injected retro-orbitally into 5–7-week-old female athymic nude mice. 16 hours post-injection, mice were sacrificed, and lungs, liver, brain, and bones were collected. EV accumulation was evaluated using the Odyssey imaging system (LI-COR Biosciences), and fluorescence signals were quantified in arbitrary units with ImageJ software by measuring NIR signal intensity across whole-organ images (García-Silva et al., 2021).

### EV-induced vascular permeability assay

To evaluate the impact of EVs on vascular permeability, we followed the protocol described by Hergueta et al., 2025 (Hergueta-Redondo et al., 2025). Briefly, 5 μg of EVs derived from scramble control or CD133 KO cells were administered via retro-orbital injection every three days over three weeks into 5– 7-week-old female athymic nude mice. 1 hour after the final EV injection, mice received a tail vein injection of fluorescently labeled dextran (1 mg/g body weight), and lung permeability was analyzed one hour later using bioluminescent imaging (Xenogen IVIS-200 Optical IVIS). To generate NIR fluorescent dextran, 6.5 mg of dextran was diluted in 1.8 mL of 0.1 M NaHCO₃. A total of 190 μL of DMSO and 10 μL of Cyanine 7.5 NHS ester (25.6 mM) were added to the solution. The reaction mixture was kept protected from light and stirred at RT for 4 hours. Labeled dextran was then purified using 50 kDa Amicon Ultra centrifugal filters (Millipore) through a two-step centrifugation protocol: 14,000 rpm for 10 minutes (filtration), followed by recovery at 7,500 rpm for 5 minutes.

### Mammary fat pad studies

To evaluate in vivo tumor growth, orthotopic injections of human scramble control and CD133 KO cells were performed in 5–7-week-old female athymic nude mice. Cells (1Å∼ 10⁵ per site) were resuspended in 25 μL of Matrigel (Corning; catalog no. 356221) mixed with 25 μL of PBS (total volume 50 μL) and kept on ice before injection. Mice were anesthetized with ketamine/xylazine (100/10 mg/kg, s.c.), and injection sites were sterilized with ethanol. A small incision was made near the fourth mammary gland, and 50 μL of cell suspension was injected into both left and right mammary fat pads using an insulin syringe. Incisions were closed with sutures, and analgesia was administered postoperatively (Temgesic 0.05–0.1 mg/kg, s.c.). Mice were monitored until recovery and maintained in sterile cages, changed every 3 days.

Eight weeks post-injection, animals were anesthetized, tumors exposed, and volumes measured with calipers. Tumors and lungs were collected and fixed in formalin for histological analysis. After 24 h, tissues were transferred to 70% ethanol, paraffin-embedded, and processed for immunohistochemistry targeting CD133 and pan-cytokeratins.

### Statistical analysis

All statistical analyses were conducted using GraphPad Prism version 10.5.0. Quantitative data are presented as mean ± standard error of the mean (SEM). For comparisons between two groups, the Student’s t-test was used. For comparisons involving more than two groups, either one-way ANOVA or the Kruskal-Wallis test was applied, depending on data distribution. Post hoc analyses included Tukey’s or Dunn’s multiple comparisons tests, as appropriate. For experiments involving two independent variables, two-way ANOVA followed by Dunnett’s multiple comparisons test was used to compare each condition to a designated control group. A p-value < 0.05 was considered statistically significant. Significance is indicated as follows: p < 0.05 (∗), p < 0.01 (∗∗), p < 0.001 (∗∗∗), and p < 0.0001 (∗∗∗).

## RESULTS

### CD133 is enriched in basal-like TNBC and correlates with poor prognosis

To assess the clinical relevance of CD133 (encoded by *PROM1*) in BC, we analyzed its expression across BC subtypes using publicly available datasets and a patient cohort from the Curie Institute previously described (Maire et al., 2013a; 2013b; Suresh et al., 2022). *PROM1*/*CD133* mRNA levels were significantly higher in TNBC compared to hormone receptor-positive (HR⁺) and HER2⁺ subtypes (Fig. S1A).

Transcriptomic profiling further revealed that, within TNBC cell lines, *PROM1*/*CD133* mRNA expression was selectively enriched in BL1 and BL2 subtypes, while only background levels were detected in MES and LAR subtypes (Fig. 1A, B). Importantly, high *PROM1*/*CD133* mRNA expression correlated with shorter recurrence-free survival in TNBC patients (Fig. S1B).

**Figure 1.**
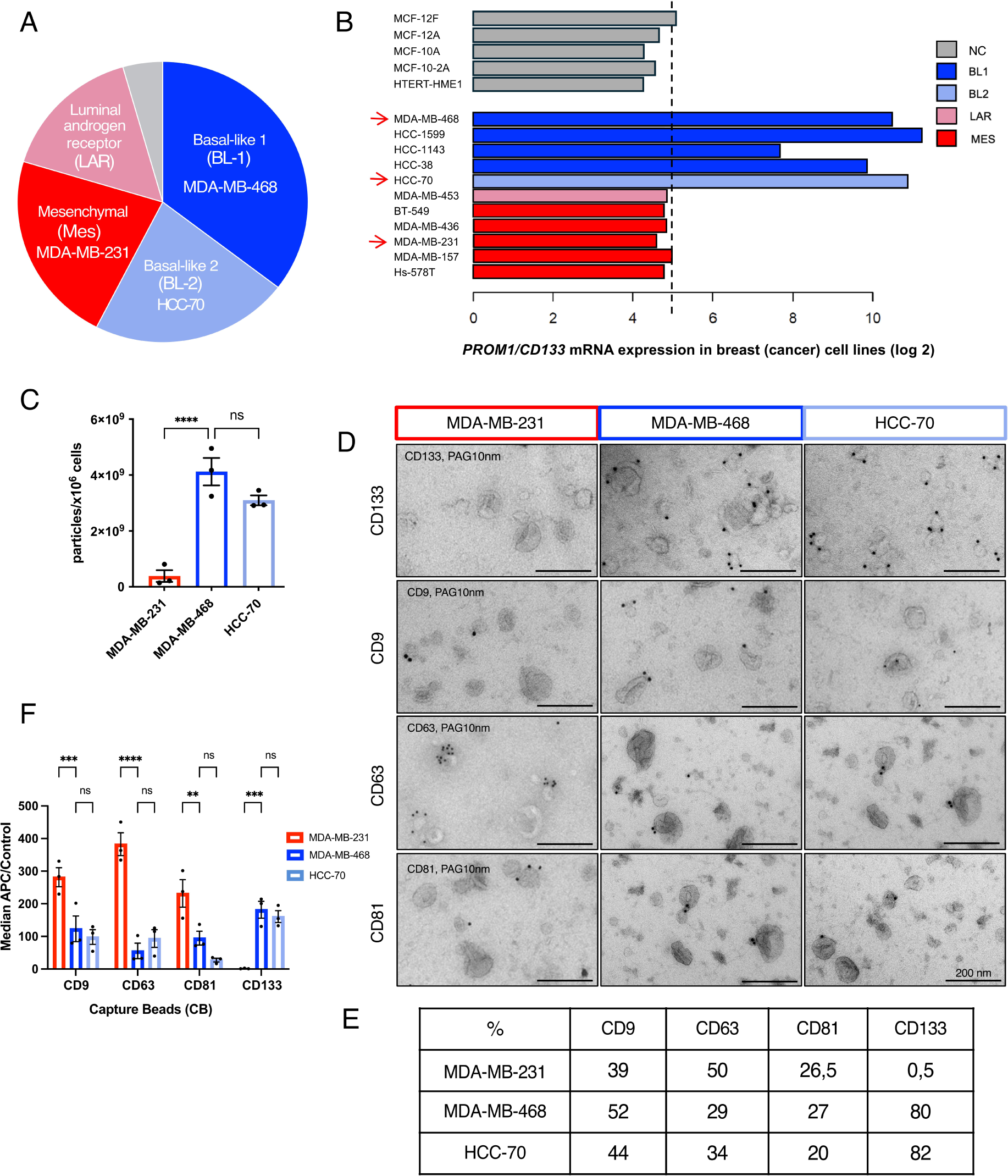
CD133 expression and EV profiling across TNBC subtypes. **A**. Schematic illustration of the distinct molecular subtypes of TNBC cells were used in this study: MDA-MB-231 (MES), MDA-MB-468 (BL-1), and HCC70 (BL-2). **B**. *PROM1/CD133* mRNA expression (log2 transformed) in various breast/ and breast cancer cell lines. NC (grey) corresponds to non-cancerous cell lines. TNBC cell lines are depicted according to the “Lehmann TNBC subtype” nomenclature: basal-like 1 (BL1, dark blue), basal-like 2 (BL2, blue-gray), luminal androgen receptor (LAR, sky blue), mesenchymal (MES, red) (Lehmann et al., 2011, 2016). Red arrows indicate the cell lines used in this study. **C**. Histogram showing EV concentration (particles per million cells) for the three TNBC cell lines, measured by nFCM. Data are presented as mean ± SEM from 3 independent experiments, analyzed by one-way ANOVA is shown. \*\*\*\**P* = < 0.0001; ns = 0.0901. **D**. Top: Representative IEM images of EVs labeled with CD133, CD63, CD9, and CD81, demonstrating marker localization in EVs from TNBC cells (MDA-MB-231, MDA-MB-468, HCC70). Data from 200 EVs per condition. Scale bar: 200 nm. **E.** Relative quantification of the percentage of positive CD133, CD63, CD9, and CD81 EVs. **F.** Multiplex bead-based FC (MACSPlex) of EV surface markers. Captured EVs from the three TNBC cell lines were detected using a mix of APC-conjugated antibodies against CD9, CD63, CD81, and CD133. MFI values were corrected using antibody-incubated beads without EVs as background. Data are shown as mean ± SEM from 3 independent experiments; analyzed by two-way ANOVA. For CD9: \*\*\**P* = 0.0007; ns = 0.7327; For CD63: ns = 0.5128; \*\*\*\**P* = < 0.0001; For CD81: ns = 0.1550; \*\**P* = 0.0027; For CD133: ns = 0.8028, \*\*\**P* = 0.0002.

These findings identify CD133 as a marker of BL-TNBC with prognostic values, delineating a functionally distinct subpopulation. Given its enrichment in basal-like TNBC and previous links between CD133 to EV secretion from plasma membrane protrusions (Marzesco et al., 2005; Hurbain et al., 2022; D’Angelo et al., 2023; Pleskač et al., 2024; Gomez-Duro et al., 2025), we hypothesized that BL-TNBC cells release a specialized population of CD133^+^ EVs consistent with PD-EVs, with CD133 potentially acting as both a marker and a regulator of their biogenesis.

### BL-TNBC cells release CD133⁺ EVs with subtype-specific marker profiles suggestive of PD-EVs

To test this hypothesis, three TNBC cell lines were selected based on their *PROM1*/*CD133* mRNA expression: MDA-MB-468 (BL1), HCC70 (BL2), both expressing high *PROM1*/*CD133* mRNA levels, and MDA-MB-231 (MES) with low or absent mRNA expression (Fig. 1B). EVs were isolated by differential ultracentrifugation (100K pellet), quantified by nanoparticle tracking analysis (NTA) and nano-flow cytometry (nFCM), and characterized by Western blot (WB) and immuno-electron microscopy (IEM), following the MISEV guidelines (Théry et al., 2018; Welsh et al., 2024).

BL1 (MDA-MB-468) and BL2 (HCC70) cells released significantly more EVs per million cells than the MES cell line MDA-MB-231. nFCM quantification showed 4×10⁹, 3×10⁹, and 4×10⁸ particles per 10⁶ cells, respectively (Fig. 1C). Despite these differences in EV abundance, vesicle size distribution was comparable across subtypes (Fig. S1C), indicating that elevated EV production in BL-TNBC cells reflects increased EV secretion rather than vesicle size differences.

WB analysis confirmed the presence of canonical EV markers (CD63, CD81, CD9, Alix, Tsg101, and Syntenin) in representative EV preparations (Fig. S1D). Notably, CD133 was abundant in EVs from BL-TNBC cells but nearly undetectable in MDA-MB-231-EVs, whereas CD63 displayed the inverse pattern: enriched in MDA-MB-231-EVs and reduced in BL-TNBC-EVs (Fig. S1D, E), consistent with subtype-specific EV biogenesis routes.

IEM confirmed vesicular identity and marker distribution, revealing ∼80-82% of BL-TNBC-EVs positive for CD133 compared to ∼0.5% in MDA-MB-231-EVs, while ∼50% of MDA-MB-231-EVs were CD63⁺ versus ∼30% in BL-TNBC-EVs (Fig. 1D, E). Multiplexed EV profiling (MACSPlex)further reinforced the subtype-specific differences, showing CD133 enrichment in BL-TNBC-EVs and CD63 predominance in MDA-MB-231-EVs (Fig. 1F).

Together, these results indicate that BL-TNBC cells release a distinct population of CD133⁺ EVs with features consistent with PD-EVs. Although both BL1 and BL2 cells released abundant CD133⁺ EVs, we focused subsequent functional studies on the BL1-TNBC model (MDA-MB-468), the most prevalent BL subtype (∼35% of TNBC) (Lehman et al., 2011 ; 2016), to test CD133-dependent PD-EV biogenesis.

### CD133 is required for efficient EV secretion and membrane organization in MDA-MB-468 cells

To directly determine whether CD133 drives the biogenesis and secretion of these EVs, we generated a CD133 knockout (CD133 KO) MDA-MB-468 cell line using CRISPR/Cas9. Sequencing confirmed indels within the *PROM1* locus, validating the knockout. This was further corroborated by RT-qPCR, which showed an ∼80% reduction in *PROM1* mRNA expression (Fig. S2A), and by flow cytometry (FC), which revealed a near-complete loss of surface CD133 (∼0.7% in KO cells versus ∼85% in MDA-MB-468 (WT) cells) (Fig. 2A). Importantly, scramble control cells displayed a CD133 surface profile comparable to WT cells (Fig. 2A).

**Figure 2.**
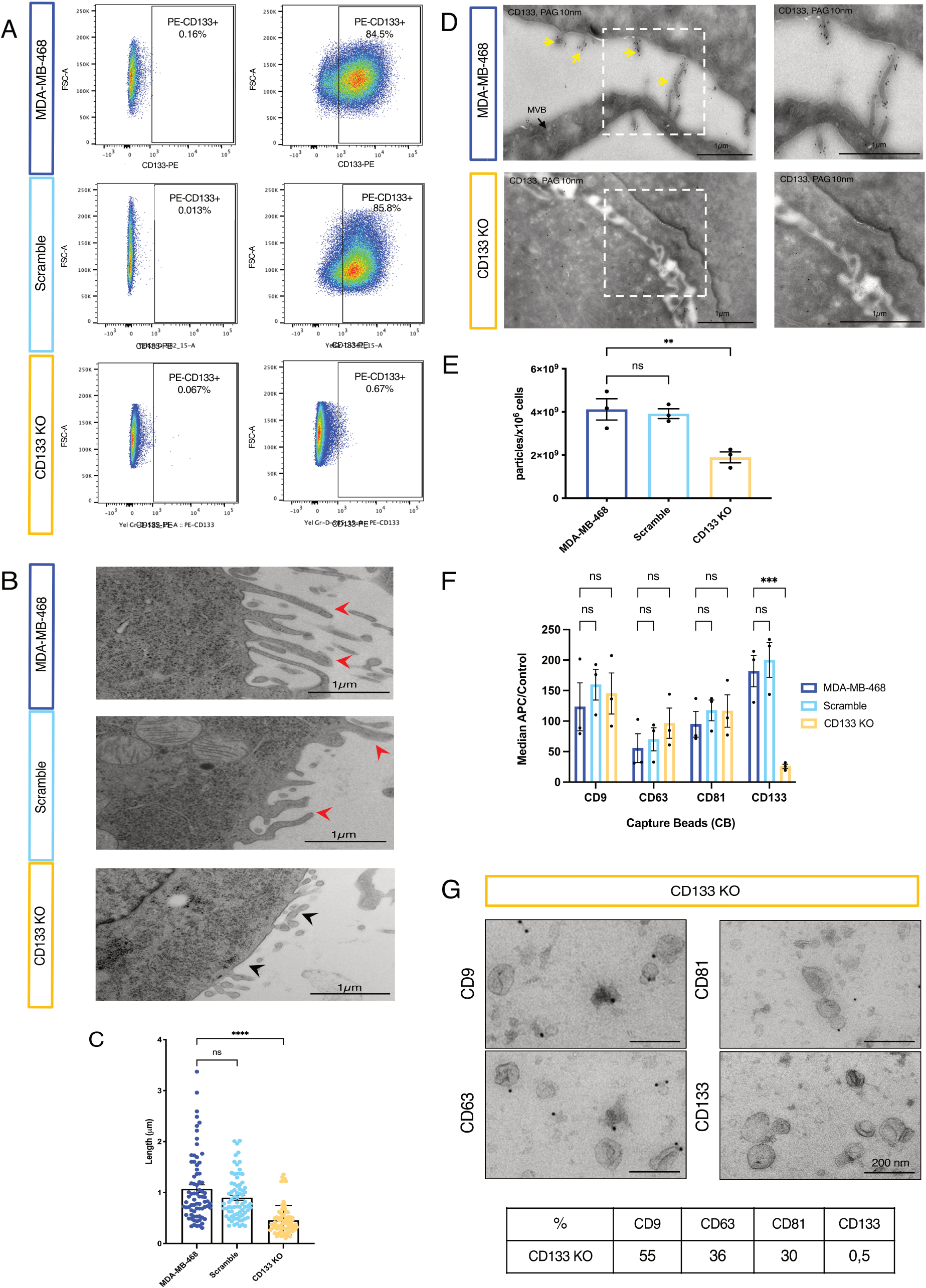
CD133 knockout reduces EV release and alters the surface marker profile of MDA-MB-468-derived EVs. **A**. FC analysis showing CD133 surface expression in MDA-MB-468 (WT), scramble control and CD133 KO cells, using a PE-conjugated anti-CD133 antibody. Gating was performed on singlet live cells (DAPI–, FSC-A vs FSC-H) before quantifying PE-positive events. The same gating strategy was applied consistently across all groups. **B.** Representative TEM micrographs of WT, scramble control, and CD133 KO cells. Red arrows indicate membrane protrusions and black arrowheads mark amorphic protrusions. Scale bar: 1 µm. **C.** Quantification of protrusion length and number (n = 75). Mean ± SEM; one-way ANOVA. ****P = < 0.0001; ns = 0.0501. **D**. Representative ultrathin cryosections of WT and CD133 KO cells immunolabeled with anti-CD133-PAG (10 nm). Yellow arrows indicate CD133⁺ staining localized in membrane protrusions. Right: magnification of the white boxed region. Scale bars are indicated in each image. **E**. Histogram showing the number of EV particles per million cells in WT, scramble control, and CD133 KO conditions quantified by nFCM. Mean ± SEM from 3 independent experiments, analyzed by one-way ANOVA. \*\**P* = 0.0071; ns = 0.8886. **F**. MACSPlex bead-based assay of EV surface marker profiling. EVs captured from WT, scramble control, and CD133 KO cells were detected using APC-conjugated antibodies against CD81, CD63, and CD9. Data are shown as mean ± SEM from 3 independent experiments, analyzed by two-way ANOVA. For CD9: ns = 0.5074 (WT vs scramble); ns = 0.7713 (WT vs CD133 KO). For CD63: ns = 0.8881 (WT vs scramble); ns = 0.4271(WT vs CD133 KO). For CD81: ns = 0.7563 (WT vs scramble); ns = 0.7750 (WT vs CD133 KO). For CD133: ns = 0.8314 (WT vs scramble); \*\*\**P* = 0.0004 (WT vs CD133 KO). **G**. Top: Representative IEM images of EVs from CD133 KO cells immunolabeled for CD9, CD63, CD81, and CD133. 200 EVs analyzed per condition. Scale bar: 200 nm. Bottom: Quantification of the percentage of EVs positive for each marker.

Next, we examined the impact of CD133 loss on cellular architecture. Transmission electron microscopy (TEM) revealed elongated, well-organized plasma membrane protrusions in WT and scramble control cells, whereas CD133 KO cells exhibited shorter and irregular extensions (Fig. 2B, C). IEM localized CD133 specifically to these protrusions in WT cells, while KO cells lacked CD133 and showed amorphous extensions (Fig. 2D). These results indicate that CD133 is required to maintain organized epithelial-like membrane architecture (Fig. 2B–D).

EV secretion was then evaluated in CD133 KO, WT and scramble control cells. nFCM quantification revealed an approximately two-fold reduction in EV release by CD133 KO cells (2×10⁹ particles per 10⁶ cells) compared to WT and scramble control cells (4.1×10⁹, and 3.9×10^9^ particles per 10⁶ cells, respectively; Fig. 2E), while vesicle size distributions remained unchanged (Fig. S2B, C).

WB analysis confirmed the marked reduction of CD133 in both EVs and cell lysates from CD133 KO cells, while canonical EV markers remained largely unaffected (Fig. S2D, E; Fig. 5C). Consistently, MACSPlex profiling of SEC-isolated EVs detected CD133 only in EVs from WT and scramble control and confirmed the expected EV marker profile (Fig. 2F).

IEM validated the vesicular identity and marker distribution, revealing that only ∼0.5% of EVs from CD133 KO cells were positive for CD133 (Fig. 2G), consistent with the very low CD133 expression in these cells (∼0.7%) (Fig. 2A) and comparable to the negligible labeling observed in MDA-MB-231 EVs (Fig. 1D, E).

Together, these results demonstrate that CD133 is functionally required for EV secretion and is incorporated into these vesicles. By sustaining organized protrusions and epithelial-like membrane architecture, CD133 contributes to EV biogenesis and secretion. Thus, CD133 acts as a key regulator of EV dynamics in MDA-MB-468 BL-TNBC cells, linking membrane organization to vesicle secretion.

### CD133 regulates the molecular cargo and signaling potential of CD133^+^ EVs in MDA-MB-468 cells

To assess whether CD133 influences not only EV secretion but also their molecular composition and signaling potential, we performed label-free quantitative proteomics (LFQ-MS) on EVs derived from WT and CD133 KO MDA-MB-468 cells. Analyses were conducted across five independent biological replicates per condition, allowing a robust comparison of CD133-EV cargo.

Unsupervised hierarchical clustering revealed tight intra-group consistency and clear segregation between conditions, indicating that CD133 depletion substantially alters EV composition (Fig. 3A). Differential abundance analysis, visualized as a volcano plot, confirmed the selective enrichment of PROM1 (CD133) in WT EVs, consistent with our previous IEM, MACSPlex, and WB results (Fig. 2F, G; S2D, E).

**Figure 3.**
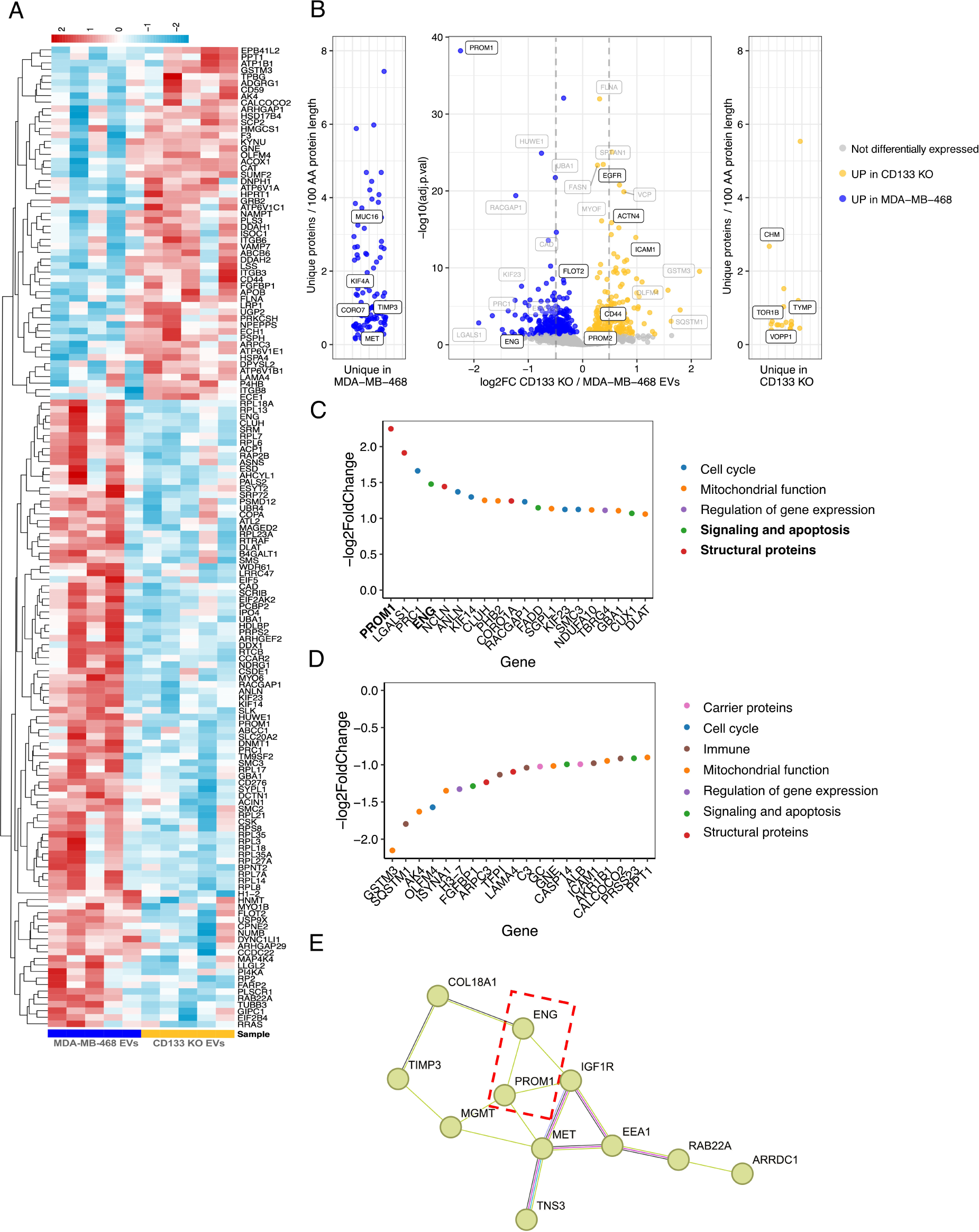
CD133 shapes the proteomic landscape of EVs derived from MDA-MB-468 and CD133 KO cells. **A**. Heatmap representing the top 150 proteins with the highest absolute mean Z-scores across all samples, selected as representative of the EV proteomic signature. Z-score normalization was applied to LFQ intensity values, and hierarchical clustering was based on Euclidean distance and single linkage. Color gradients indicate relative protein abundance, ranging from –2 (low) to +2 (high). **B**. Volcano plot illustrating global proteomic differences between EVs from WT and CD133 KO cells (n = 5 biological replicates per condition). log₂FC is plotted against –log₁₀ (adjusted p-value). Proteins with a |log₂FC| ≥ 1 and adjusted p-value ≤ 0.05 are considered significantly differentially expressed. **C-D**. GO functional classification of the 20 proteins with the highest (C) and lowest (D) abundance ratios in EVs from WT versus CD133 KO cells. Proteins are color-coded by functional category, with PROM1 (CD133-related) and ENG (CD105-related) highlighted in bold. **E**. PPI network generated using the STRING database, centered on CD133 and its associated proteins. The network highlights potential interaction partners and enriched functional pathways in WT-derived EVs. The red dashed rectangle outlines a subnetwork containing PROM1(CD133), ENG (endoglin/CD105).

Beyond CD133, proteins enriched in CD133^+^ EVs, included KIF23 and PRC1 (mitotic spindle regulators) and ENG (Endoglin/CD105), a pro-angiogenic TGF-β co-receptor (Dallas et al., 2008) (Fig 3B). Gene ontology (GO) enrichment revealed that CD133^+^ EVs from WT cells were enriched in proteins involved in cell cycle progression, mitochondrial bioenergetics, and apoptotic signaling, whereas CD133 KO EVs contained proteins linked to metabolic detoxification, immune response, and ECM remodeling (Fig. 3C, D). A protein–protein interaction (PPI) network further highlighted a PROM1-ENG (CD133-CD105) cluster among proteins enriched in CD133⁺ EVs (Fig. 3E).

These data indicate that CD133 not only drives EV biogenesis but also shapes their molecular composition. CD133^+^ EVs carry proteins linked to cell cycle regulation, mitochondrial function, apoptotic signaling, and angiogenesis, highlighting their potential to mediate tumor-stroma communication in BL-TNBC. Given their enrichment in vesicles released from protrusion-rich cells, these EVs may correspond to PD-EVs, a possibility further explored in subsequent functional analyses.

### CD133^+^ EVs promote angiogenesis independently of EV uptake or endothelial proliferation

Building on our proteomic data showing CD133-dependent enrichment of angiogenesis-related proteins in EVs, we next evaluated their functional impact on endothelial cells. Human umbilical vein endothelial cells (HUVECs) treated with CD133⁺ EVs from WT cells displayed significantly enhanced capillary-like tube formation on Matrigel, characterized by increased network complexity and branching compared to cells treated with CD133 KO EVs or PBS. Notably, the angiogenic response induced by CD133⁺ EVs was at least comparable to, and in some cases exceeded that elicited by VEGF (Fig. 4B, C), underscoring their potent pro-angiogenic potential.

**Figure 4.**
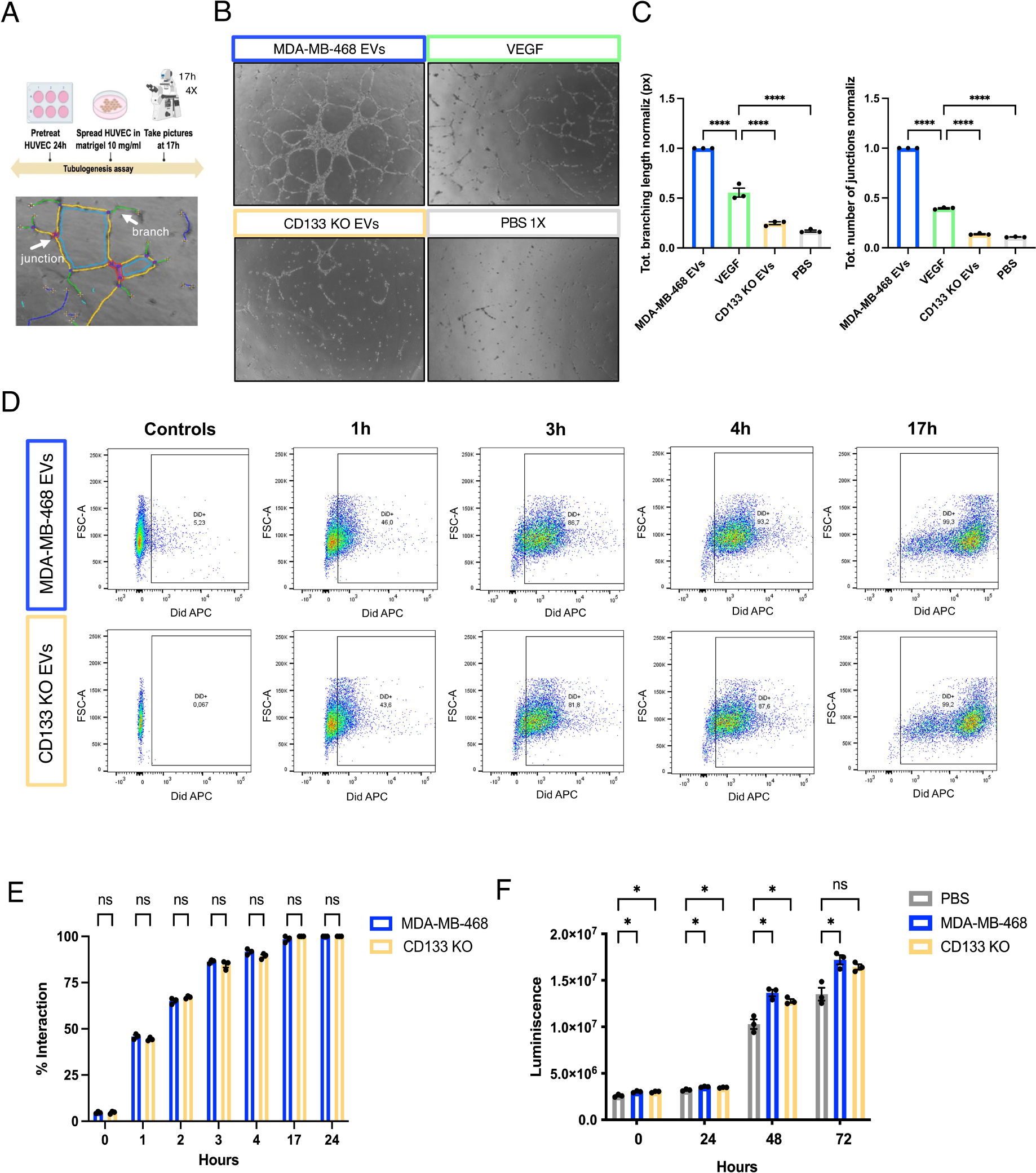
CD133^+^ EVs enhance tubulogenesis independently of uptake or proliferation of endothelial cells. **A**. Schematic of the experimental workflow and macro-assisted analysis using ImageJ. Quantified parameters are shown: total number of junctions and total branching length. **B**. Representative images of capillary-like tube formation by HUVECs 17 hours post-treatment with EVs from WT or CD133 KO cells (40X magnification). **C.** Quantification of tube formation (number of junctions and total branching length). Data are shown as mean ± SEM from 3 independent experiments; two-way ANOVA. ****P < 0.0001. **D**. Uptake of DiD-labeled EVs by HUVECs at different time points, assessed by FC. No qualitative differences between WT and CD133 KO EV uptake were observed. **E**. Quantification of DiD-labeled EVs uptake by HUVECs at different times. Data are shown as mean ± SEM from 3 independent experiments. Comparisons between WT and CD133 KO EVs at each time point were analyzed by unpaired t-test. At 0 h ns = 0.9999; at 1 h ns = 0.7323; at 2 h ns = 0.3529; at 3 h ns = 0.4569; at 4 h ns = 0.1806; at 17 h ns = 0.7102; at 24h ns > 0.9999. **F**. Endothelial cell proliferation assay following EV treatment, showing similar mitogenic effects for both WT and CD133 KO EVs. Data are shown as mean ± SEM from 3 independent experiments; one-way ANOVA. At 0 h \**P* = 0,0242 (PBS vs WT) and \**P* = 0.0271 (PBS vs CD133 KO). At 24 h \**P* = 0.0215 (PBS vs WT) and \**P* = 0.0474 (PBS vs CD133 KO). At 48 h \**P* = 0.0133 (PBS vs WT) and \**P* = 0.0422 (PBS vs CD133 KO). At 72 h \**P* = 0.0258 (PBS vs WT) and ns = 0.599 (PBS vs CD133 KO).

To determine whether this effect reflected differences in EV internalization, we tracked the uptake of DiD-labeled EVs by HUVECs using FC. Both CD133⁺ and CD133 KO EVs displayed similar internalization kinetics, with ∼90% of cells positive at 4 hours and near-complete uptake by 17 hours (Fig. 4D, E), indicating that CD133 does not affect the efficiency of EV uptake by endothelial cells.

We next asked whether enhanced angiogenic activity was driven by endothelial proliferation. Both CD133⁺ and CD133 KO EVs modestly stimulated proliferation relative to PBS controls, but their effects were undistinguishable (Fig. 4F), ruling out proliferation as the primary driver of the angiogenic phenotype.

Together, these findings demonstrate that the pro-angiogenic activity of CD133⁺ EVs is not due to differences in EV internalization or mitogenic signaling. Instead, it likely reflects the selective enrichment of bioactive cargo in a CD133-dependent manner. These results support a model in which CD133 influences cargo composition and angiogenic potential by y linking membrane organization to vesicle function. While our data point to a role for CD133 in shaping EV cargo with features of PD-EVs, the precise mechanisms underlying this process remain to be determined.

### CD133^+^ EVs exhibit organ-specific accumulation

To complement our *in vitro* findings on PD-EV-mediated angiogenesis, we next asked whether CD133^+^ EVs influence organ-specific biodistribution *in vivo.* We first assessed whether CD133 expression affects tumor-intrinsic behavior. Orthotopic injection of scramble control or CD133 KO cells into the mammary fat pad of immunocompromised mice revealed comparable tumor volumes and growth kinetics over 60 days (Fig. 5A). At endpoint, CD133 expression was maintained in parental tumors but absent in CD133 KO-derived tumors, validating the sustained loss of CD133. Immunohistochemistry (IHC) for pan-cytokeratins showed no significant differences in epithelial differentiation or tumor composition between groups (Fig. 5B, C), indicating that CD133 expression does not alter primary tumor growth or epithelial differentiation.

**Figure 5.**
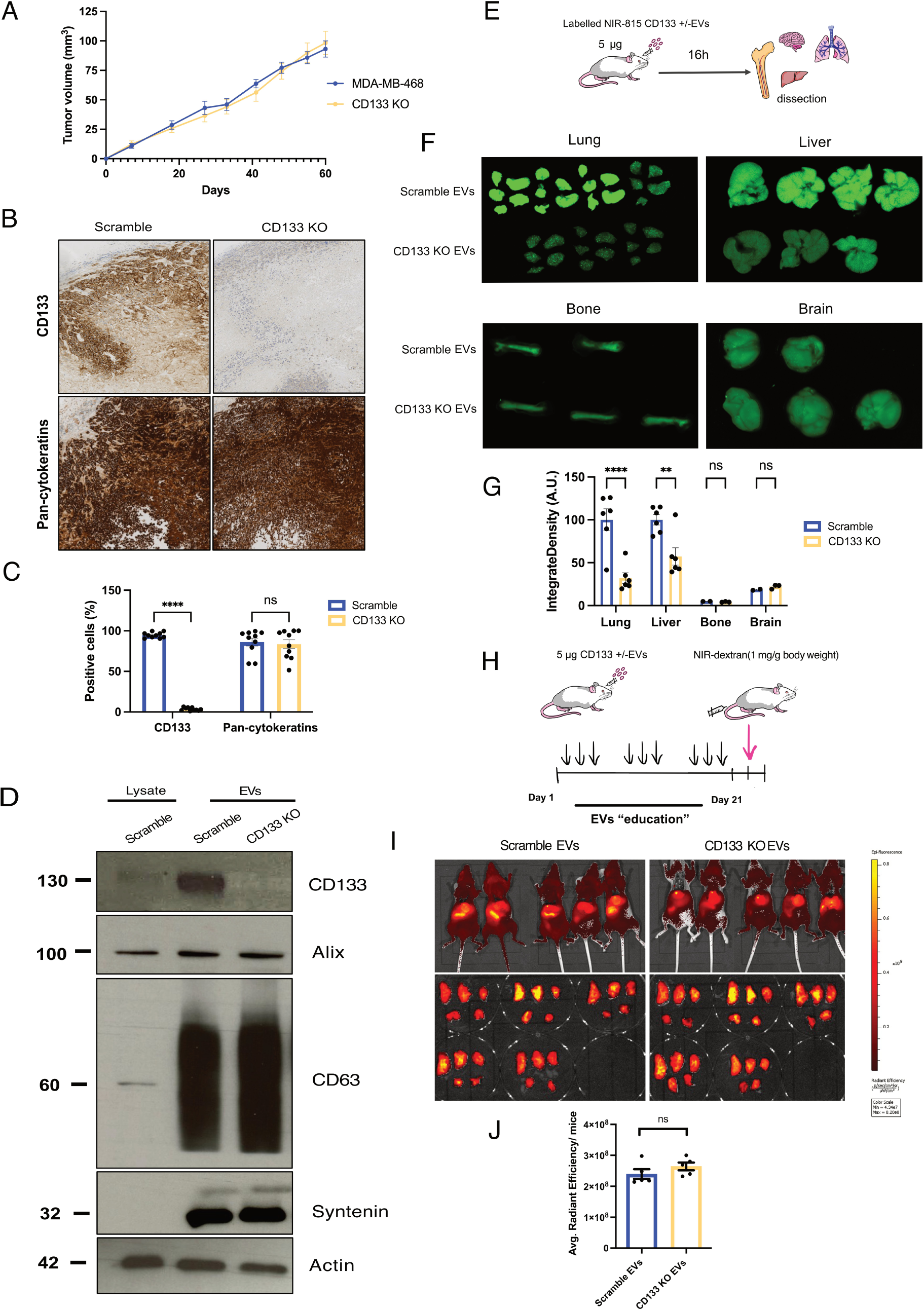
Functional impact of CD133^+^ EVs on cancer progression and organotropic dissemination. **A**. Tumor growth curves over 60 days following orthotopic injection of scramble control or CD133 KO MDA-MB-468 cells into the mammary fat pad. Data are presented as mean ± SEM from one experiment (n = 5 mice/group). **B**. Representative IHC images of endpoint tumors stained for CD133 and pan-cytokeratins. **C**. Quantification of IHC staining. CD133 expression was retained in tumors derived from parental cells and absent in CD133 KO tumors; no significant differences were observed in epithelial identity . Data are shown as mean ± SEM from one experiment; n = 5 mice/group; two-way ANOVA. ns = 0.6102; \*\*\*\**P* < 0.0001. **D**. WB analysis of whole-cell lysates (20 µg) and EVs (20 µg) from scramble control and CD133 KO MDA-MB-468 cell lines, normalized by total protein input. **E**. Schematic of the *in vivo* biodistribution assay. Mice were intravenously injected with 5 µg of NIR-815 labeled CD133^+^ or CD133 KO EVs and sacrificed 16 hours later for tissue dissection. **F**. Representative NIR fluorescence images showing EVs in the lungs, liver, bone, and brain, and associated quantification. **G**. Quantification of EV biodistributiion. Data are shown as mean ± SEM from 2 independent experiments; n = 4 mice/group; two-way ANOVA. \*\*\*\**P* < 0.0001 (lung); \*\**P* = 0.0010 (liver); ns = 0.9717 (bone); ns = 0.8444 (brain). **H**. Schematic of the *in vivo* “EV education” experimental design. Mice received repeated injections of CD133^+^ or CD133 KO EVs over 21 days. **I**. Images of Texas Red-labeled dextran extravasion in lungs of CD133^+^ or CD133 KO EV-educated mice and quantification of vascular permeability. **J**. Quantification of vascular permeability. No significant differences were observed between groups. Data are shown as mean ± SEM from one experiment; n = 5 mice/group; unpaired t-test. ns = 0.2222.

Given that tumor cell-intrinsic behavior remained unchanged, we next asked whether CD133⁺ EVs influence metastatic niche conditioning, in line with prior reports that tumor-derived EVs drive organ-specific metastasis (Peinado et al., 2012; Hoshino et al., 2015). Near-infrared-labeled EVs from scramble control and CD133 KO cells were retro-orbitally injected into athymic nude mice (Fig. 5D, E). Sixteen hours post-injection, CD133^+^ EVs preferentially accumulated in the lung (∼50% of total signal) and liver (∼35%), whereas CD133 KO EVs showed significantly reduced localization to these sites. No differences were observed in brain or bone (Fig. 5F, G).

To evaluate whether organ-specific tropism was linked to vascular permeability, mice were preconditioned with repeated injections of CD133⁺ or CD133 KO EVs (Fig. 5H) and pulmonary endothelial leakiness was assessed by dextran extravasation. No significant differences were detected between the groups, indicating that organ-specific accumulation occurs independently of vascular permeability changes (Fig. 5I, J).

Together with our *in vitro* data, these results demonstrate that CD133 shapes EV biodistribution and functional activity, consistent with a role in niche priming, while acting independently of tumor cell-intrinsic growth properties. This model decouples EV-dependent effects from tumor-intrinsic behaviors, supporting the central role of CD133 in orchestrating EV function.

### CD105 is enriched in CD133^+^ EVs

Proteomic profiling revealed selective enrichment of CD105 in CD133⁺ EVs from WT cells, prompting us to investigate its potential contribution to their pro-angiogenic effects (Dallas et al., 2008; Tian et al., 2018). Consistent with the proteomic data, CD105 levels were markedly reduced in EVs from CD133 KO cells, as confirmed by WB and MACSPlex analysis (Fig. 6A, B; Fig. S3A).

**Figure 6.**
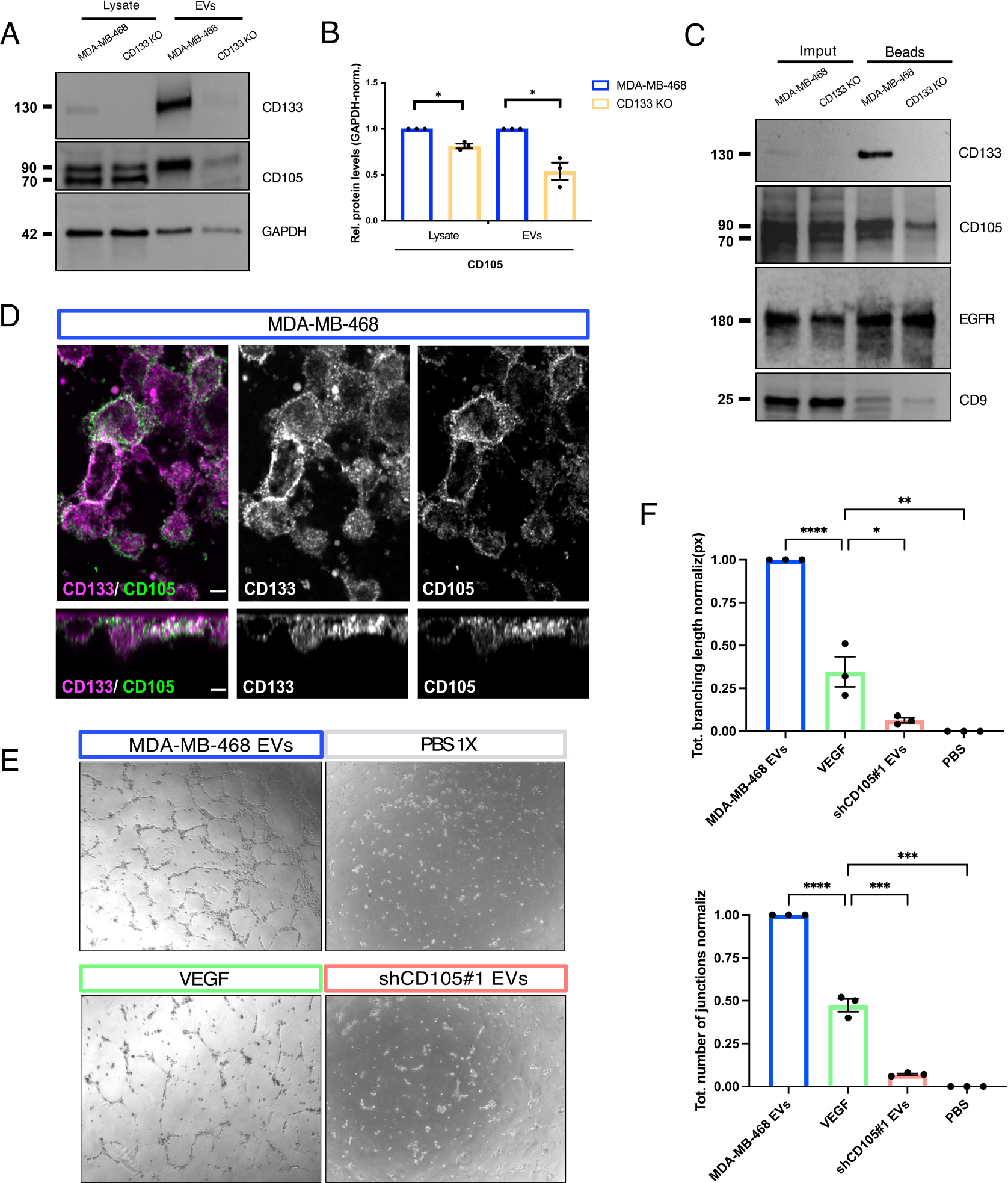
CD133 and CD105 co-localize at the cell surface and are enriched in EVs with angiogenic functions. **A**. WB analysis of whole-cell lysates (12 µg) and EVs (1 × 10⁹ particles) from WT or CD133 KO cells confirms the presence of both proteins in EVs and their modulation upon CD133 loss. **B**. Quantification of CD105 protein levels in cell lysates and EVs normalized to GAPDH. Data are shown as mean ± SEM from 3 independent experiments; unpaired t-test. \*\**P* = 0.0058; \*\*\**P* = 0.0003, respectively. **C**. Cell surface biotinylation assay showing membrane localization and co-enrichment of CD133 and CD105 at the plasma membrane in WT and CD133 KO cell lines (n = 3 independent experiments). **D**. Single confocal xy and orthogonal z-sections images of WT cells stained for CD133 (magenta) and CD105 (green). Scale bar: 5 µm. **E**. Representative images of capillary tube formation assays (17 h, 40X magnification) following EV treatment from WT or shCD105 KD cells (n = 3 independent experiments). **F**. Quantitative analysis of junction number and branch lengths. Note the impaired angiogenic potential of EVs upon CD105 knockdown. Data are represented as mean ± SEM of 3 independent experiments; two-way ANOVA. For branching length: \*\*\*\**P* < 0.0001; \**P* = 0.0163; \*\**P* = 0.0046. For number of junctions: \*\*\*\**P* < 0.0001; \*\*\**P* = 0.0005 (VEGF vs shCD105#1); \*\*\**P* = 0.0002 (VEGF vs PBS).

At the transcriptional level, *ENG* mRNA was elevated in several BL and selected MES-TNBC cell lines, but not in MDA-MB-231 cells (Fig. S3B). This expression pattern closely paralleled that of *PROM1* (Fig. 1B), suggesting possible co-regulation or functional interplay between CD133 and CD105 in EV-producing cells. Although CD105 has not previously been implicated in EV biogenesis, STRING-based PPI network and text-mining analyses further support a potential cooperative role in EV formation or selective cargo loading, possibly in coordination with CD133 (Fig. 3E).

### CD133 and CD105 localize at the plasma membrane and in lipid raft microdomains

CD133 binds to cholesterol, partitions into cholesterol-rich lipid raft microdomains and interacts with the actin cytoskeleton to support membrane protrusions and EV budding from protrusion-rich regions (Röper et al., 1997; Corbeil et al., 2001; Giebel et al., 2004; Karbanová et al., 2018; Thamm et al., 2019; Hurbain et al., 2022). Given the selective enrichment of CD105 in CD133^+^ EVs (Fig. 3B, C), we hypothesized that CD133 may facilitate CD105 recruitment to the plasma membrane and its incorporation into EVs released from protrusions.

To test this, surface biotinylation under non-permeabilizing conditions showed a significant reduction of CD105 at the plasma membrane in CD133 KO cells (Fig. 6C), suggesting that CD133 is required for CD105 membrane localization or retention. Membrane-associated CD9, a tetraspanin involved in EV biogenesis (Duke et al., 2025), was also decreased, whereas EGFR levels remained unchanged and served as a loading control.

Immunofluorescence (IF) further supported these findings. In MDA-MB-468 WT cells, CD133 and CD105 were observed in discrete, often adjacent membrane domains, suggesting spatial proximity without complete co-localization (Fig. 6D). CD9 and CD81 were similarly enriched in CD133⁺ membrane regions, implicating these domains as potential sites of EV formation (Fig. S3C).

Detergent-resistant membrane (DRM) fractionation provided additional mechanistic insight. In the parental MDA-MB-468 cells, CD133, CD105, and lipid raft markers (Flotillin-1, CD81) (Bickel et al., 1997; Cherukuri et al., 2004) were enriched in low-density fractions (notably fraction 3) (Fig. S3D). In CD133 KO cells, Flotillin-1 was reduced and CD81 was lost from these fractions, and CD105 remained detectable but with at lower levels, indicating impaired raft integrity. These findings indicate that CD133 is critical for CD105 retention at the plasma membrane and for maintaining raft integrity, a prerequisite for CD105 incorporation into EVs from protrusion-rich regions.

### CD105-enriched EVs from CD133^+^ cells promote angiogenesis

We next asked whether CD133-dependent recruitment of CD105 has functional consequences for the pro-angiogenic activity of CD133⁺ EVs. To test this, WT cells were transduced with control shRNA (shCtrl) or two independent shRNAs targeting CD105 (shCD105#1, #2). The most effective knockdown (shCD105#1) significantly reduced CD105 protein levels in both cell lysates and CD133^+^EVs (Fig. S3E-G).

Normalized CD133⁺ EVs (6 µg/mL total protein) were applied to HUVECs, and tube formation assays were performed. EVs from shCtrl cells significantly enhanced tubulogenesis, whereas EVs from CD105 knockdown cells (shCD105#1) showed markedly diminished pro-angiogenic activity (Fig. 6E, F). These results indicate that CD105 is functionally required for the pro-angiogenic effects of CD133⁺ EVs. The data further suggest a coordinated mechanism in which CD133 modulates cargo selection and angiogenic signaling, at least in part by promoting CD105 surface retention and lipid raft association, thereby enabling its incorporation into EVs.

## DISCUSSION

Our study demonstrates that CD133 is not merely a passive marker of cancer stem cells but an active organizer of EV biogenesis and cargo selection in BL-TNBC. By sustaining membrane protrusions and lipid raft organization, CD133 governs both the secretion and molecular composition of a distinct vesicle population. Although CD133⁺ EVs fall within the exosome size range, (i) CD133 is detected on plasma membrane protrusions by TEM/IEM, (ii) CD133 and CD105 co-fractionate with lipid raft markers by DRM fractionation, and (iii) CD133⁺ vesicles are enriched in EV isolates from protrusion-rich cells. Together, these findings support a plasma membrane origin consistent with PD-EVs, rather than endosome-derived exosomes (Marzesco et al., 2005; Hurbain et al., 2022; D’Angelo et al., 2023; Pleskač et al., 2024).

Mechanistically, CD133 is selectively enriched in cholesterol-dependent lipid raft microdomains restricted to plasma membrane (Karbanová et al., 2018; Pleskač et al., 2024), where it co-enriches with CD9, flotillin, and CD105. Knockout of CD133 led to fewer or less stable membrane protrusions, impaired PD-EV secretion, and reduced the membrane-and raft-associated pool of CD105, and supporting a model in which CD133 organizes protrusion-associated cholesterol-dependent microdomains to coordinate cargo selection and vesicle budding. These observations extend prior studies linking CD133 to microvillus remodeling and EV release from cholesterol-rich domains (Marzesco et al., 2005; Hurbain et al., 2022), positioning CD133 at the center of a subtype-specific EV biogenesis pathway.

A central finding of this study is the identification of a functional CD133-CD105 axis. CD105, a TGF-β co-receptor essential for angiogenesis (Dallas et al., 2008; Tian et al., 2018) is selectively enriched in CD133⁺ EVs. Both proteins are found in close proximity at the plasma membrane and within raft fractions, suggesting that cargo sorting occurs within spatially organized microdomains. Depletion of CD105 abolished the pro-angiogenic effects of these EVs without affecting vesicle uptake, supporting a model in which CD133 acts as a molecular scaffold that recruits CD105 via raft-associated sorting mechanisms. This challenges the view that EV cargo is stochastically incorporated and supports an emerging model of organized, membrane-based cargo selection.

Our findings support and extend the concept of “prominosomes”, CD133-marked EVs originating from cholesterol-rich apical membrane domains, independent of the endosomal route (Karbanová et al., 2025). Here, CD133 not only serves as a marker of these structures but also actively organizes lipid raft subdomains to coordinate the selective recruitment of functionally relevant cargo such as CD105. This model is reinforced by analogous systems, including ectosome formation from lipid rafts in monocytes (Del Conde et al., 2005), tetraspanin-enriched membrane compartmentalization (Charrin et al., 2003), and raft-mediated miRNA delivery (Zhou et al., 2014), supporting the notion that raft-localized sorting mechanisms dictate EV identity and function. Within this framework, the CD133-CD105 axis defines a subtype-specific EV subclass with angiogenic properties shaped by spatially regulated cargo inclusion.

Functionally, CD133⁺ EVs with PD-EV features exhibit potent pro-angiogenic activity in 3D endothelial branching assays. This effect is abrogated by CD105 knockdown, yet uptake of the vesicles by HUVECs remained intact, demonstrating that the loss of function stems from altered cargo rather than impaired delivery. In contrast, HUVEC proliferation in 2D monolayer cultures is not significantly influenced, indicating that EV-mediated signaling is context-dependent and preferentially acts on specialized endothelial subpopulations, such as tip cells. These findings emphasize that EV function is determined more by cargo composition than by delivery efficiency, consistent with models of selective cargo recruitment driven by lipid microdomains and protein-protein interactions (van Niel et al., 2018). Moreover, they preferentially accumulated in the lung and liver, a tropism not attributable to passive vascular permeability, suggesting that CD133 contributes to organotropic EV behavior and potentially to PMN formation.

Beyond angiogenesis, proteomic analysis revealed enrichment of proteins involved in cell cycle regulation, mitochondrial activity, and apoptosis in CD133⁺ EVs, suggesting additional roles in tumor survival and adaptation within the TME. The close spatial proximity of CD133 and CD105 at the plasma membrane and within lipid raft microdomains supports the CD133-CD105 axis as a central mechanism shaping angiogenic signaling. While our current data demonstrate proximity and clustering, further studies are needed to determine whether these proteins functionally interact within these EVs and how this might influence tumor aggressiveness and PMN formation.

This proximity also raises mechanistic questions. Both proteins are glycosylated (Corbeil et al. 2010 ; Fonsatti & Maio, 2004), and additional post-translational modifications, including palmitoylation, may influence their recruitment to lipid rafts and incorporation into PD-EVs (Levental et al., 2010). It remains unclear whether their spatial distribution is direct or mediated by raft-associated adaptors such as flotillins, tetraspanins, or annexins (Yáñez-Mó et al., 2009; Wang et al., 2013; Karbanová et al., 2017). Future studies will be required to investigate these possibilities and clarify how such mechanisms contribute to selective cargo sorting and PD-EV function.

Importantly, we demonstrate that EV biogenesis and function are subtype-specific. BL-TNBC cells produce abundant CD133⁺ EVs enriched with pro-angiogenic cargo, whereas MES TNBCs with low or no CD133 expression lack detectable PD-EV production. This strong association between molecular subtype and vesicle identity suggests that PD-EV biogenesis is governed by intrinsic molecular programs and aligns with previous subtype-based stratifications (Burstein et al., 2015; Lehmann et al., 2016).

Our findings also have clear translational implications. Given the ongoing clinical exploration of CD105 in angiogenesis-driven cancers (Douglas et al., 2021; Zhang et al., 2021), the enrichment of CD105 in CD133^+^ EVs underscores the potential of targeting this EV subclass for therapy. Beyond therapy, CD133⁺ EVs may serve as biomarkers for aggressive BL-TNBC and as vehicles whose disruption could impair early metastatic niche formation.

In conclusion, this study identifies a membrane-based mechanism of EV biogenesis in BL-TNBC, orchestrated by CD133 and characterized by selective CD105 incorporation. We define a CD133-CD105 axis that shapes the identity and function of a subtype-specific EV subclass with pro-angiogenic properties. By repositioning CD133 from a passive CSC marker to an active organizer of vesicle composition, our findings directly link membrane architecture to tumor-stroma communication, offering both mechanistic insight and translational opportunities.

## ACKNOWLEDGMENTS

We thank Paula Cambronera for her valuable assistance with the *in vivo* experiments, and Alix Zhou for his contribution to the development of the CD133 knockout cell line using CRISPR-Cas9. We are also grateful to Nicolas Ansart from the CurieCoretech Extracellular Vesicle Platform at Institut Curie for support with the MACSPlex assays, as well as the Flow Cytometry Platform at Institut Curie for technical assistance. We further acknowledge Dr. Yago Arribas De Sandoval for help in generating the proteomics figures in R and Dr. Lucas Alves Tavares for support with molecular biology experiments. The LSMP thanks Patrick Poullet from the bioinformatics Core facility (CUBIC) of the Institut Curie U1331 for the continuous development of myProMS and Michael Ricard from the LSMP for statistical support. This work was supported by the Centre National de la Recherche Scientifique (CNRS) and Institut Curie, Fondation ARC pour la Recherche sur le Cancer (PJA2024070008456 to G.D.), La Ligue Contre le Cancer (RS25/75-81to G.D.), the European Union, EVCA Twining project (Horizon GA no. 101079264), and the European Union’s Horizon 2020 research and innovation programme under the Marie Skłodowska-Curie (Grant agreement No 847718). M.G.D. was supported by a PhD fellowship from Institut Curie EuReCa PhD Programme (Grant agreement No 847718).

## AUTHOR CONTRIBUTION

M.G.-D., P.B., S.S.-R., M.H.-R., F.D., performed the experiments, M.G.-D., P.B., M.H.-R, Y.M., D.L., T.D, designed the experiments and analyzed the data. G.D., L.M.-J., H.P., T.D., supervised the experiments. G.D. and M.G.-D wrote the paper. G.R. provided fundings. G.D. conceived the hypothesis, oversaw the project, supervised all the work and obtained funding. All authors have read the final version of the manuscript and agree to its publication.

## CONFLICTS OF INTERESTS

The authors declare no conflicts of interest

## Data Availability Statement

All data generated or analyzed during this study are available in the article and its online supplemental material. Requests for materials should be addressed to G. D’Angelo.

## Notes

### Competing Interest Statement

The authors have declared no competing interest.

