## Supplementary figures S1-S3 for "CD133 shapes extracellular vesicle cargo and angiogenic function in basal-like triple-negative breast cancer"

**Supplementary Figure 1.**

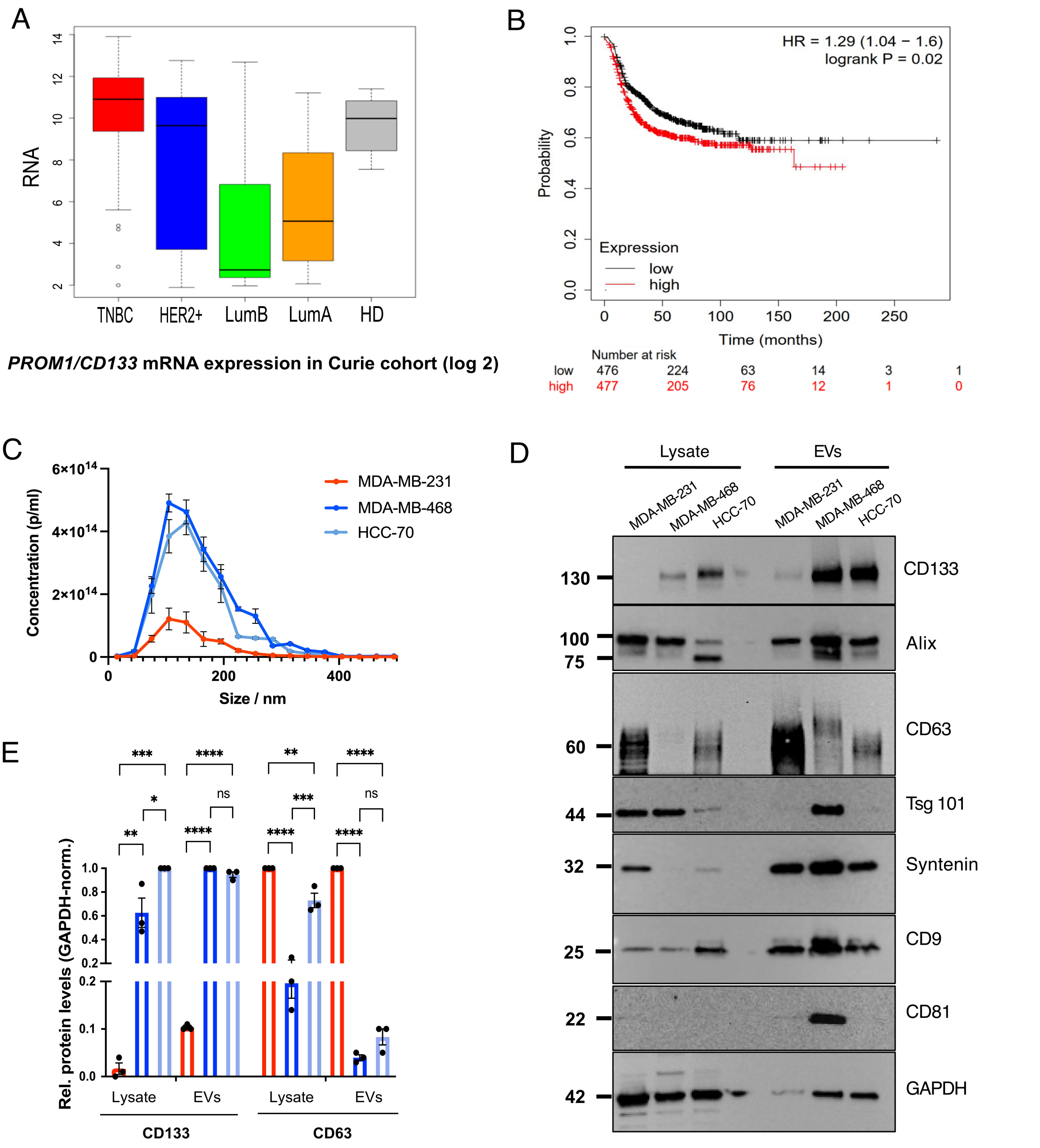


**CD133 expression in TNBC patient samples and cell lines, and characterization of EVs from TNBC cells. A**. Relative *PROM1/CD133* mRNA expression in TNBC (red), HER2 (blue), luminal B (LumB, green), luminal A (LumA, orange), and normal breast tissues (HD, grey) in the Curie cohort is illustrated by box plots ($log_2$ transformed). Outliers are shown within each population (open circles). Student's t test was used to compare RNA levels between two groups. The p values are indicated (*P < 0.05; **P < 0.01; ***P < 0.001; ns P > 0.05). **B**. High *PROM1/CD133* mRNA expression is associated with a poorer prognosis in TNBC. Recurrence Free Survival (RFS) data based on *PROM1/CD133* mRNA expression (high in red *vs*. low in black) obtained from the Kaplan–Meier (KM) plotter website (http://kmplot.com) in the TNBC population. The obtained Hazard Ratio (HR) with 95% confidence interval and log-rank p-values ($p$ = 0.02) are shown. **C**. NTA showing EV size distribution (n = 3 independent experiments) from the three TNBCCs (MDA-MB-231, MDA-MB-468, HCC70). **D**. WB analysis of whole-cell lysates (12 μg) and EVs (1 × $10^9$ particles) derived from MDA-MB-231, MDA-MB-468 (WT), and HCC70 cells, probed for CD133 and canonical EV markers. **E**. Quantification of CD133 and CD63 protein levels in cell lysates and EVs normalized to GAPDH. Data are presented as mean ± SEM from 3 independent experiments; analyzed by one-way ANOVA. In cell lysates for CD133: **$P$ = 0.0072; *$P$ = 0.0207; ***$P$ = 0.0002. In EVs for CD133: ****$P$ = < 0.0001; ns = 0.3120. In cell lysates for CD63: ****$P$ = < 0.0001; ***$P$ = 0.00024; **$P$ = 0.0096. In EVs for CD63: ****$P$ = < 0.0001; ns = 0.2710.

**Supplementary Figure 2.**

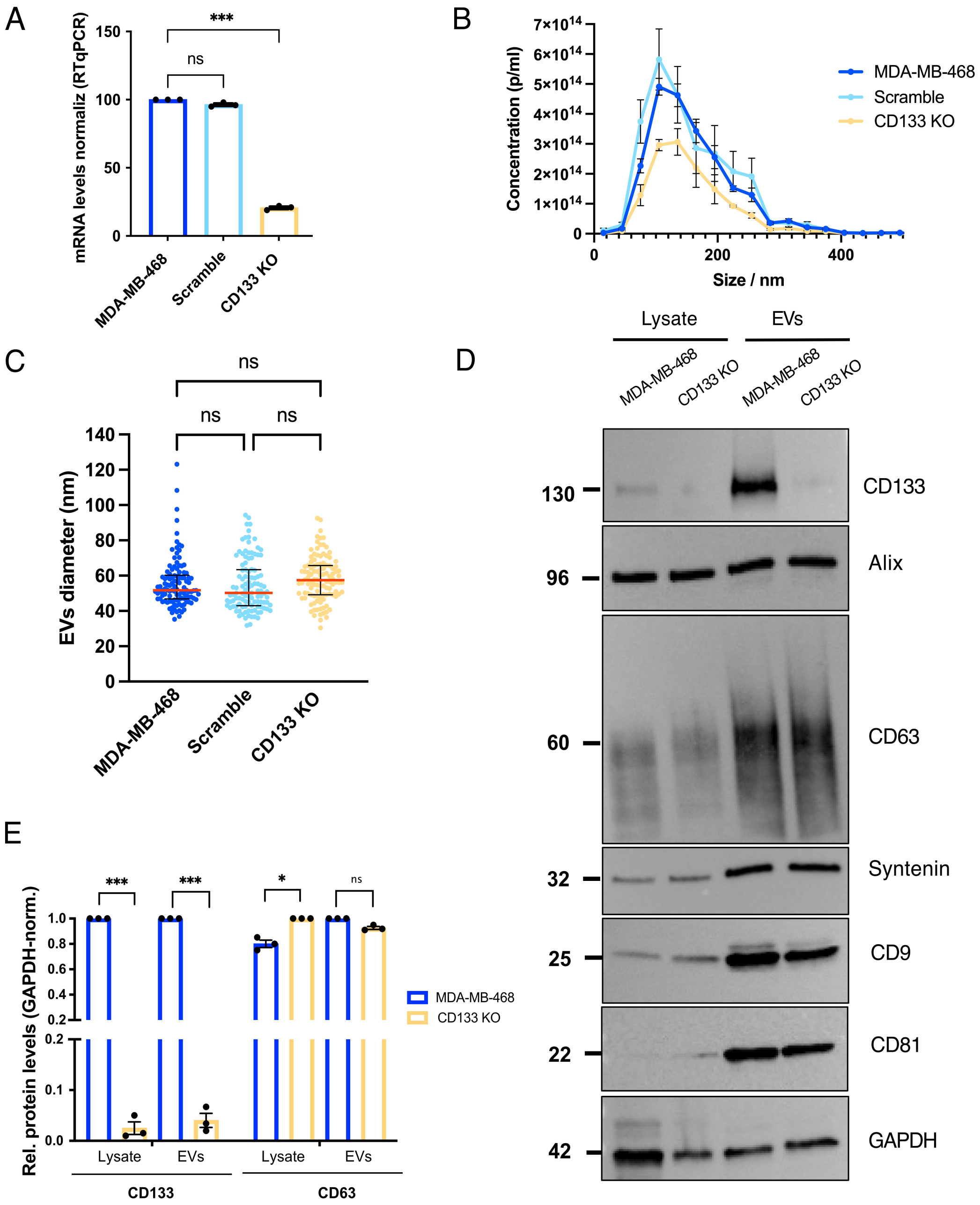


**Characterization of MDA-MB-468 cells and EVs following CD133 depletion via CRISPR-Cas9.** **A**. RT-qPCR analysis of *CD133/PROM1* mRNA expression in WT, scramble control, and CD133 KO cells, confirming effective knockout in CD133 KO cells. Mean ± SEM; one-way ANOVA. ***$P$ = 0.0003; ns = 0.1154. **B**. NTA of EV size distribution from WT, scramble control, and CD133 KO cells (n = 3 independent experiments). **C**. Quantification of EV diameter from microscopy images of WT, scramble control, and CD133 KO cells. Data from 200 EVs; mean ± SEM; one-way ANOVA. ns = 0.4098 (WT vs scramble); ns = 0.8134 (scramble vs CD133 KO); ns = 0.1430 (WT vs CD133 KO) (n = 110 per condition). **D**. WB analysis of whole-cell lysates (12 µg) and EVs ($1 \times 10^9$ particles) from WT and CD133 KO cells probed with CD133 and canonical EV markers. **E**. Quantification of CD133 and

CD63 protein levels in cell lysates and EVs normalized to GAPDH. Data are presented as mean ± SEM from 3 independent experiments; unpaired t-test. For CD133, in cell lysates ***$P$ = 0.0002 and EVs ***$P$ = 0.00023. For CD63, in cell lysates *$P$ = 0.0207 and EVs ns = 0.250.

**Supplementary Figure 3.**

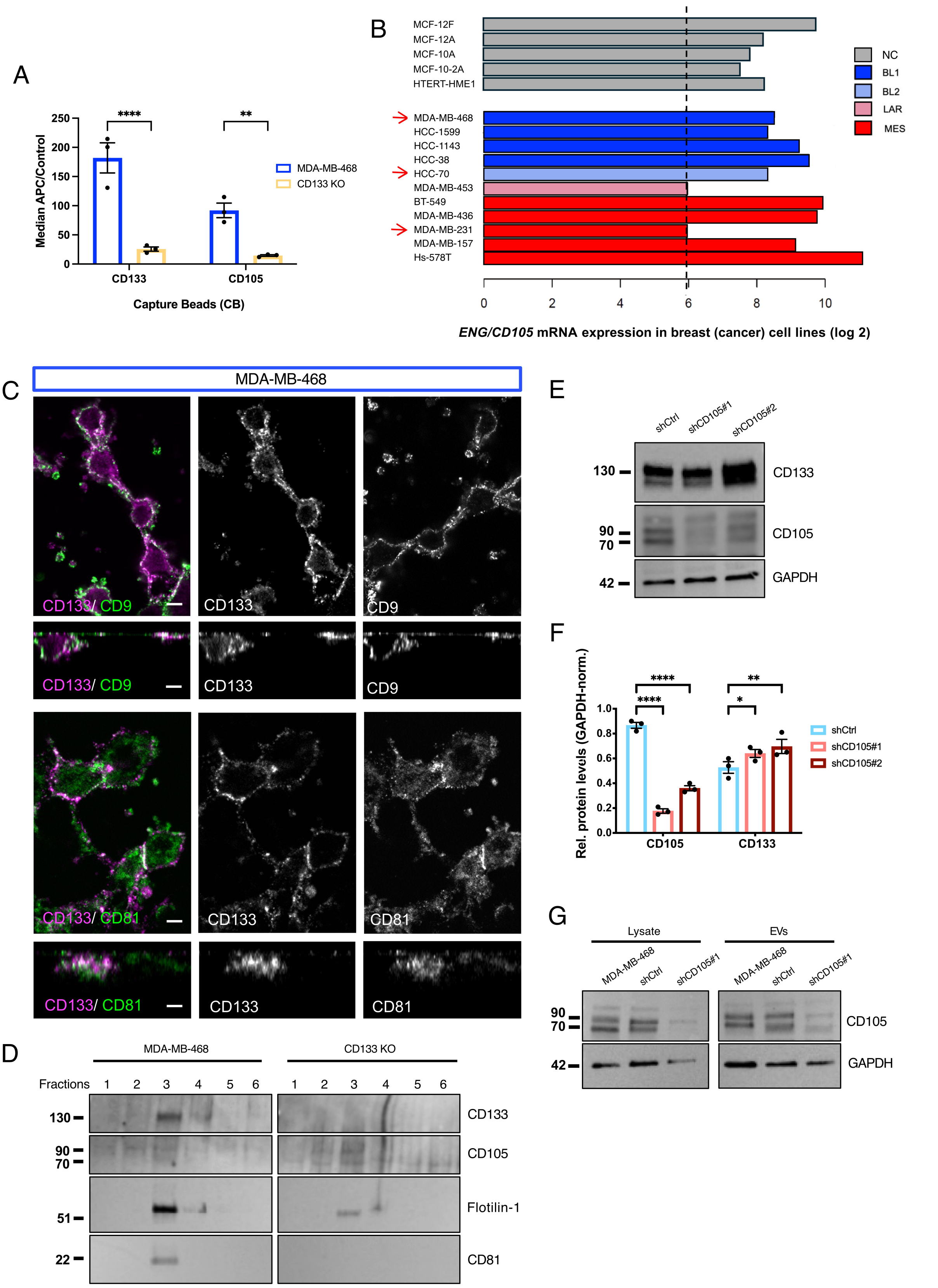


**Molecular and functional interplay between CD133 and CD105 in TNBC cells and their EVs. A.** MACSPlex analysis detecting CD133 and CD105 on EVs isolated from WT and CD133 KO cells using

APC-labelled anti-CD81, anti-CD63, and anti-CD9 antibodies. Data represent mean ± SEM for 3 independent experiments; two-way ANOVA. For CD133: ****$P < 0.0001$; ns = 0.6862. For CD105: *$P = 0.0157$; ns = 0.9582. **B**. *ENG/CD105* mRNA expression (log2 transformed) in various breast/ and breast cancer cell lines. NC (grey) corresponds to non-cancerous cell lines. TNBC cell lines are depicted according to the "Lehmann TNBC subtype" nomenclature: basal-like 1 (BL1, dark blue), basal-like 2 (BL2, blue-gray), luminal androgen receptor (LAR, sky blue), mesenchymal (MES, red) (Lehmann et al., 2011, 2016). Red arrows indicate the cell lines used in this study. **C**. Representative single confocal xy and orthogonal z-sections images of WT cells stained for CD133 (magenta; grey) and CD9/CD81 (green; grey). Scale bar: 5 µm. **D**. Lipid raft fractionation assay showing the partitioning of CD133 and CD105 into DRM fractions (n = 3 independent experiments). **E-F**. WB analysis and associated quantification of whole-cell lysates from WT cells transduced with scramble control shRNA (shCtrl) or two independent CD105-targeting shRNAs (shCD105#1 and shCD105#2), showing efficient knockdown. Data represent mean ± SEM for 3 independent experiments; one-way ANOVA. For CD105 ****$P < 0.0001$. For CD133 *$P = 0.0121$; **$P = 0.0011$, respectively. **G**. WB analysis of whole-cell lysates and EVs from WT transduced with shCtrl or shCD105#1, showing reduced CD105 levels and altered EV cargo upon CD105 knockdown.
